# Single-cell cytometry via multiplexed fluorescence prediction by label-free reflectance microscopy

**DOI:** 10.1101/2020.07.31.231613

**Authors:** Shiyi Cheng, Sipei Fu, Yumi Mun Kim, Weiye Song, Yunzhe Li, Yujia Xue, Ji Yi, Lei Tian

**Affiliations:** Department of Electrical and Computer Engineering, Boston University, Massachusetts 02215, USA; Department of Biology, Boston University, Massachusetts 02215, USA; Department of Philosophy & Neuroscience, Boston University, Massachusetts 02215, USA; Department of Biomedical Engineering, Boston University, Massachusetts 02215, USA; Department of Medicine, Boston University School of Medicine, Boston Medical Center, Massachusetts 02118, USA; Department of Biomedical Engineering and Ophthalmology, Johns Hopkins University, Baltimore 21231, USA

**Author notes:** These authors contributed equally to this work (Shiyi Cheng, Sipei Fu, Yumi Mun Kim).

## Abstract

Traditional imaging cytometry uses fluorescence markers to identify specific structures, but is limited in throughput by the labeling process. Here we develop a label-free technique that alleviates the physical staining and provides highly multiplexed readouts via a deep learning-augmented digital labeling method. We leverage the rich structural information and superior sensitivity in reflectance microscopy and show that digital labeling predicts highly accurate subcellular features after training on immunofluorescence images. We demonstrate up to 3× improvement in the prediction accuracy over the state-of-the-art. Beyond fluorescence prediction, we demonstrate that single-cell level structural phenotypes of cell cycles are correctly reproduced by the digital multiplexed images, including Golgi twins, Golgi haze during mitosis and DNA synthesis. We further show that the multiplexed readouts enable accurate multi-parametric single-cell profiling across a large cell population. Our method can dramatically improve the throughput for imaging cytometry toward applications for phenotyping, pathology, and high-content screening.

## Introduction

Cell morphology features are powerful phenotypical readouts, which have been the basis for pathology for decades. They are also the underlying mechanisms for varieties of imaging cytometry and high-content screening platforms to characterize pathological changes and responses to drug treatments (*1*). The most widely used approach for imaging readouts are fluorescence labels that highlight specific subcellular components or cell functions through immunofluorescence (IF), fluorescent reporter cells or dyes. However, the throughput of these approaches is fundamentally limited by the physical process of labeling. The IF staining is labor-intensive and generally requires cell fixation that does not allow kinetic observations of live cells over time. Fluorescent reporter cells are permissive for longitudinal live cell imaging. However, the process of gene editing and validation takes a significant amount of time and can be difficult to introduce multiple markers within the same cells for multiplexed analysis. Regardless of the fluorescence labeling approaches, the overlapping of the fluorescence emission spectra further limits the multiplexing capability. To alleviate these limitations, here we develop a label-free single-cell cytometry that is highly multiplexed and can forgo the physical staining via a deep learning (DL)-augmented digital labeling method.

Our work relies on the premise that label-free scattering-based microscopy captures rich structural information and can be effective to characterize cell morphological features (*2*). Brightfield, phase contrast and differential interference contrast (DIC) microscopy have been routinely used for observing and quantifying cell morphology (*3*). Scattering-based microscopy and tomography techniques have been increasingly utilized to reconstruct cellular structures (*4*). Of particular interest is the reflectance-mode microscopy that provides exquisite sensitivity in detecting nanoscale structural changes beyond the diffraction limit (*5–7*). By capturing backscattering signals, reflectance imaging provides access to the highest spatial-frequency components in the reciprocal Fourier space and thus can provide higher structural contrast than the transmission techniques (*2*). Indeed, our recent work shows that backscattering signals allow resolving finer details than the transmission counterparts (*8, 9*). In this work, we further leverage the higher sensitivity provided by the reflectance-mode microscopy and demonstrate how enriched label-free information allows predicting highly accurate subcellular structural features.

The framework of this study is summarized in Fig. 1. The angle-dependent backscattering features are captured with darkfield oblique illumination and paired with IF images (Fig. 1A). Using the IF images as the ground truths, multiple DL models are independently trained for individual IF labels (Fig. 1B). Once all the models are trained, we perform digital multiplexing by feeding the same label-free input to each network and make different IF predictions in parallel (Fig. 1C). By doing so, multiple subcellular structures and cell states can be revealed simultaneously without physical labeling. While previous works have shown that DL models can disentangle the complex structures captured in the label-free data and make *in-silico* fluorescence labeling with high accuracy (*10–12*) or holistically capture “hidden” structural features that are not easily perceived or described (*13–22*), these results are fundamentally limited by the weak structural contrast from the transmission modes that contain only forward scattering information. By exploiting the enhanced resolution and sensitivity in the backscattering data, we demonstrate a dramatic increase in the fluorescence prediction accuracy with up to 3× improvement as compared to the current state-of-the-art.

**Fig. 1.**
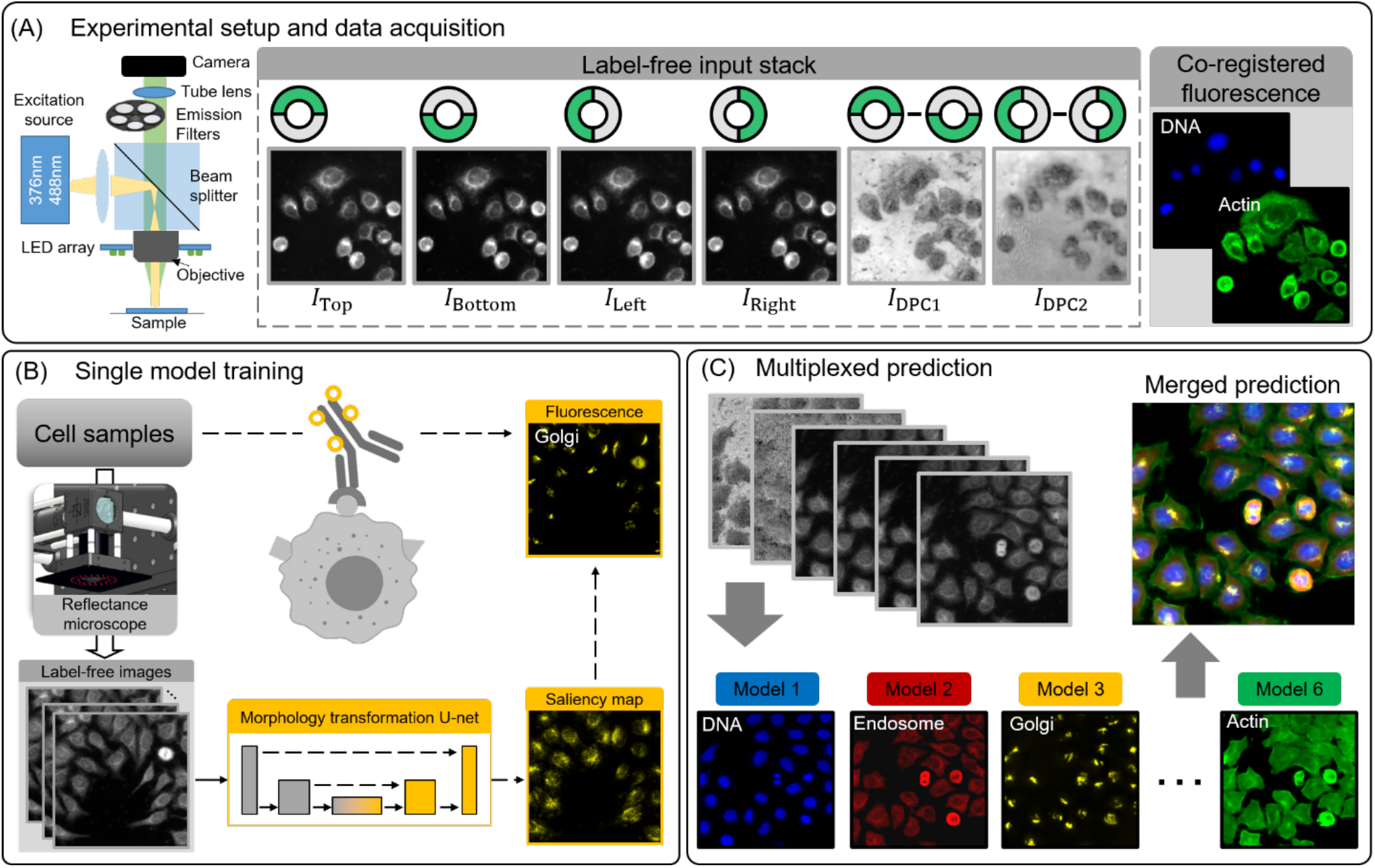
Overview of the DL augmented label-free cytometry technique. **(A)** A multimodal LED-array reflectance microscope is developed to acquire co-registered label-free reflectance and fluorescence images. Reflectance images from oblique darkfield illumination and computed drDPC contain rich morphological information and are the multi-channel input to our DL model. Two-channel epi-fluorescence images are acquired on the same sample to serve as the ground truth for training our DL model. **(B)** Individual DL models are trained independently with paired label-free and IF images. The saliency map is used to reveal specific label-free features captured by the model to perform the transformation. **(C)** To perform digital multiplexed predictions, the same reflectance input is fed to each network and makes six different IF predictions in parallel.

One distinct contribution of our work is to advance beyond the prediction of fluorescence images, and to demonstrate accurate structural phenotyping and quantitative single-cell cytometry using digitally multiplexed fluorescence images. Importantly, we show that our DL model can correctly capture and predict characteristic subcellular features during the cell cycle, including morphological changes of nuclei and Golgi apparatus. We also show that our DL model can capture the structural features of cell proliferation and recapitulate the DNA duplication through the cell cycle. Remarkably, the label-free structural features identified in proliferating cells are not obvious by visual inspection of the raw images, demonstrating that our holistic DL model can potentially capture novel cellular attributes with high accuracy. Another distinct advantage of multiplexing several fluorescence markers is that it enables the development of multiple quantitative metrics for imaging cytometry and phenotyping across the large cell population and at the single-cell level (*e.g.* cell size, nuclear-cytoplasmic ratio, nuclear roughness, Golgi eccentricity, *etc.*). As other “omic” platform technologies are rapidly being developed to evaluate biological processes at various levels (e.g. genomics, transcriptomics, proteomics, metabolomics), single-cell level structural metrics will complement these population-based studies, particularly when a contextual phenotypic shift is expected only for a subset of cells. As a demonstration, we evaluate several cellular features, including morphology and fluorescence expressing intensity on the DL predicted digitally multiplexed readouts.

A common criticism of DL-based methods is the “black-box” nature of these models (*23*). To overcome this issue, we adapt the attention mechanism (*24*) to elucidate on the working mechanism of our DL model. We construct the saliency map that highlights the most important subcellular features contributing to each IF prediction by the network. Our results show that the structural components in label-free reflectance input that correspond to the fluorescence labels can be correctly identified by the saliency map, and the “attention” is consistent across different cell batches when predicting all IF labels. This indicates that our network learns to extract the *salient* and *specific* structural information from the reflectance images matching the underlying subcellular components. In addition, the improved prediction accuracy is attributed to the enhanced resolution and sensitivity to subcellular structures from the backscattering information.

## Results

### Oblique illumination-based reflectance microscopy captures rich morphological information

The imaging platform is based on our recently developed LED-array reflectance microscope for capturing co-registered label-free reflectance and fluorescence images (*8*). By flexibly controlling the LED patterns, this new platform enables capturing multiple angle-dependent backscattering contrasts in the darkfield without any mechanical switching. Based on our prior work (*8*), we heuristically optimize the illumination strategy and implement half-annulus LED patterns along four different orientations (including top, bottom, left, and right) (see Fig. 1A). In addition, we compute the darkfield reflectance differential phase contrast (drDPC) based on the raw measurements (see Materials and Methods). The raw oblique-illumination darkfield and drDPC images contain complementary structural contrasts. In particular, subcellular structures are shown with high contrast in the raw darkfield measurements, including the nuclei, nucleoli, and hyper-reflective structures at the nuclear periphery. Cell membranes with sharp boundaries are highlighted in the drDPC images, with cytoplasm spreading on the substrate with thicker nuclei at the cells’ centers.

The extra cell topography information exhibited in the drDPC images are found particularly useful for predicting IF labels that are sensitive to morphology of the cell boundary. These label-free images are used as the multi-channel input to our DL model. On the same platform, two-channel epi-fluorescence images are concurrently acquired on the same sample to serve as the ground-truth for training our DL models (Fig. 1A) (see Materials and Methods). The significance of this new microscopy platform is that we capture enriched label-free information by multiple contrasts in the reflectance-mode. This empowers our label-free high-content cytometry technique to uncover highly sensitive and specific structural phenotypes at the single-cell level across large cell populations.

### Individual fluorescence prediction achieves state-of-the-art performance

To evaluate the performance of our DL models, we take measurements on fixed HeLa cells containing in-total six IF labels, including DNA (Hoechst), Golgi apparatus (GM130), endosome (EEA1), actin (Phalloidin), proliferation (EdU), and apoptosis (TUNEL). Specifically, five separate batches of IF staining are performed with GM130, EEA1, Phalloidin, EdU, TUNEL, each of which is co-stained with Hoechst (see Materials and Methods). We then train six networks for performing individual IF label predictions using paired reflectance-fluorescence image dataset (Fig. 1B). Additional details about the network implementation and the data preprocessing procedure are provided in Materials and Methods and Supplementary Materials Figs. S1 and S2, respectively.

A major goal of our study is to investigate the structural contrast captured by different label-free reflectance modes and understand how they impact the DL-based fluorescence predictions. To this end, we conduct ablation studies on the drDPC input in Supplementary Materials Fig. S3 and Table S1. As expected, the additional drDPC channels substantially improve the actin IF label predictions by ~9% since drDPC clearly highlights cell topography and boundaries as compared to the plain darkfield images. Notably, the additional drDPC channels also dramatically improve the prediction accuracy for proliferation and apoptosis IF labels, by ~8% and ~17%, respectively. We hypothesize that this is because distinct structural features exhibited during proliferation or apoptosis can be more prominently displayed by drDPC and subsequently recognized by the DL model.

After training, we first evaluate each network’s prediction accuracy on unseen reflectance input from the same cell batch. Figure 2 shows the label-free input, the individual IF groundtruth, and the prediction for all six labels. The predicted subcellular structures and cell states have excellent visual agreement with the ground-truths. Characteristic morphological features are clearly recovered, including rounded nuclei, cytoplasmic endosome, spreading cell membrane (actin), and Golgi apparatus at the nuclei periphery. Selective cellular events or functions such as proliferation and apoptosis are also captured by the DL predictions. Additional examples of the prediction results are shown in Supplementary Materials Fig. S4.

**Fig. 2.**
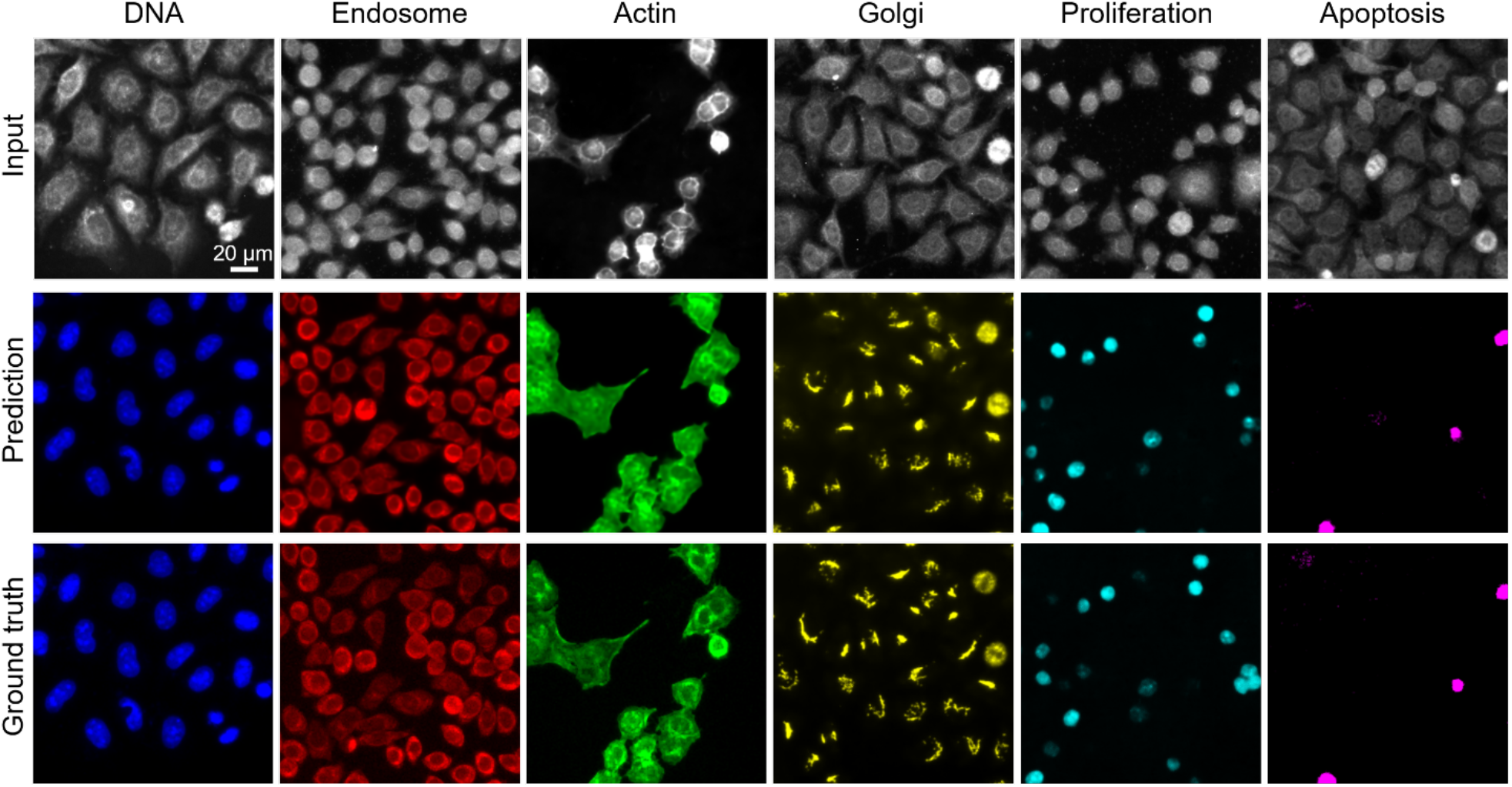
Visualization of the results from the six IF prediction networks. The rows show sample darkfield reflectance images from each input stack, the network’s IF prediction, and the groundtruth IF image, respectively. The columns show six IF labels covering four different subcellular features, including nuclei (DNA), endosome, actin, and Golgi apparatus, as well as two different cell states, including proliferation and apoptosis. The IF predictions have excellent visual agreement with the ground-truths in all six cases.

We quantify the prediction accuracy by computing several evaluation metrics on the network’s predictions based on the underlying cytometry tasks. Specifically, we formulate the predictions of the DNA, endosome, actin, and Golgi apparatus as regression problems because they are distinct subcellular structures and thus better described in their morphologies. Accordingly, the performance is quantified by calculating the image patchwise Pearson correlation coefficient (PCC), which measures the pixel-level similarity of morphologies between the DL prediction and the ground truth (see Materials and Methods). The distributions of the PCCs for the four IF label predictions are shown in the violin plots in Fig. 3A. Notably, all four of our regression DL models achieve higher accuracy as compared to the current state-of-the-art techniques based on transmission label-free microscopy data (*10*). The median PCCs on the DNA, endosome, actin, and Golgi apparatus label predictions achieve 87.25%, 91.85%, 92.01%, and 59.82%, respectively. These results agree well with the example visualizations in Fig. 2. The cellular features in the reflectance images associated with the corresponding fluorescence label are clearly visible. Although these scattering signals are entangled with other signals in the raw label-free images, our result shows that our DL models are able to recognize and distill these salient features with high accuracy. As visualized in the violin plots and quantified by the 25% and 75% quantiles in Fig. 3A, different amounts of variations in the prediction accuracy are seen for the four labels. In general, we observe that the larger deviation of the prediction from the groundtruth, as measured by the median value, is also associated with larger accuracy variations. To investigate these spatial variations, we visualize the patch-wise PCC as a spatial map in Supplementary Materials Fig. S5. Low-value PCC outliers are generally observed in background regions, and hence are removed by a standard algorithm (see Materials and Methods). Other than the background outliers, the PCCs are consistent across all the cell regions for all the four IF label predictions, which demonstrates the overall robustness of the DL model.

**Fig. 3.**
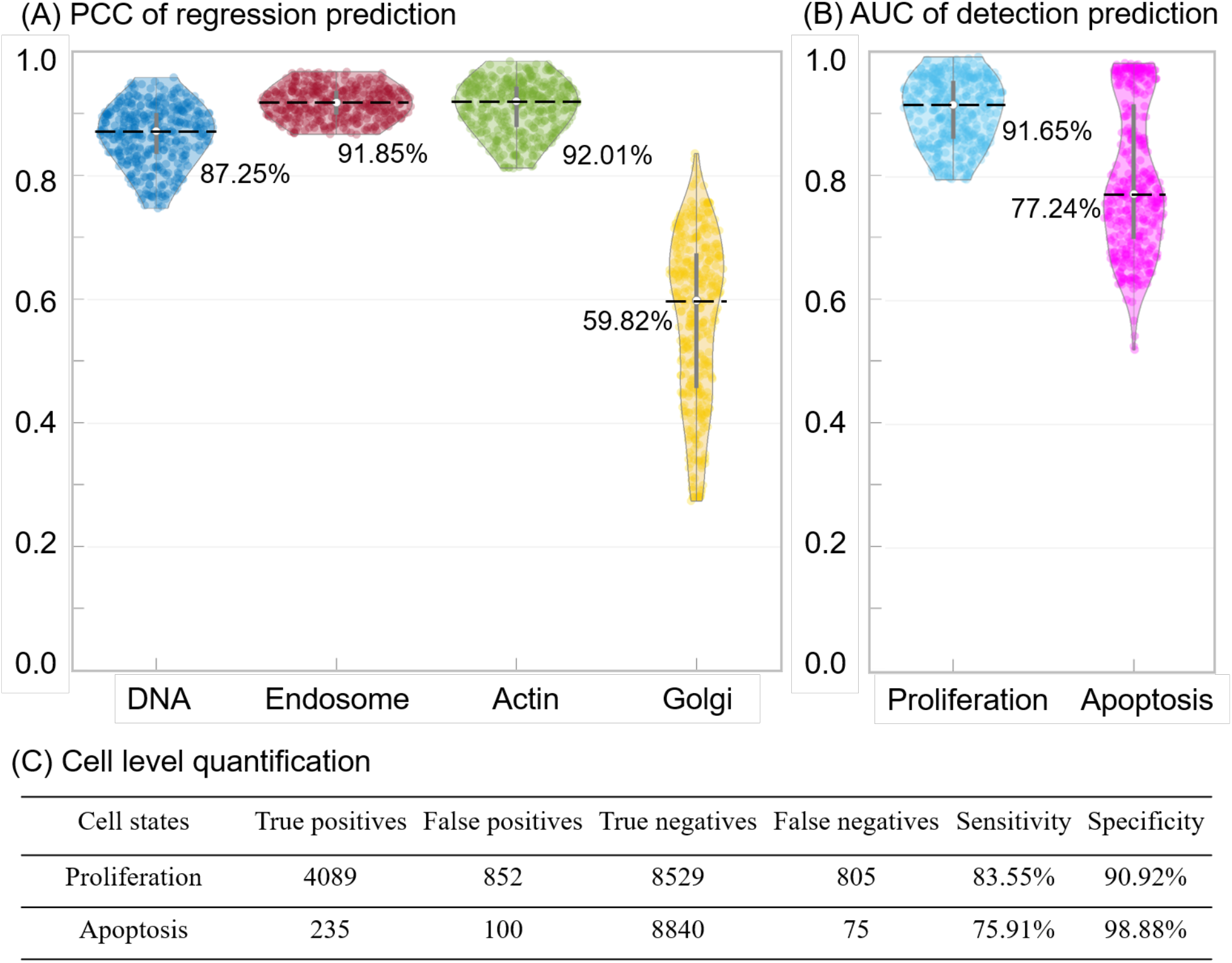
Quantitative evaluation of the DL prediction. The violin plots show the quantitative metrics for each IF label prediction. The upper and lower bounds of each grey bar represent the 25% and 75% quantiles, respectively; the center white point marked by the black dashed line denotes the median. In total, 676 testing image patches are aggregated for computing the statistics for each label. **(A)** The image patch-wise PCCs of the predictions for nuclei (DNA), endosome, actin, and Golgi apparatus, which evaluates the pixel-level similarity between the regression-type predictions and ground-truth subcellular features. **(B)** The image patch-wise AUC of the proliferation and apoptosis predictions, which assess the pixel-level detection accuracy. **(C)** Quantitative evaluation of the cell-level detection performance of the proliferation and apoptosis predictions.

We formulate the prediction of the proliferation and apoptosis as detection problems, which is more biologically meaningful because they are selective cell states. The proliferation labels are only present in the DNA-replicating cells. The apoptosis labels are only for cells undergoing programmed deaths. Accordingly, the performance is first quantified by the pixel-level Area Under the Receiver Operating Characteristic (ROC) Curve (AUC) to measure the ability of separating the positives (*i.e.* those expressing the fluorescence) from the negatives at each pixel (see Materials and Methods). The calculation is image batchwise, the same way as we calculated PCC. The distributions of the AUCs over all batches for the two IF labels are shown in the violin plots in Fig. 3B. The median detection accuracies are 91.65% and 77.24% for the proliferation and apoptosis, respectively. Next, we further evaluate the cell-level detection performance to give a more direct assessment on these two label predictions. To do so, we perform single-cell segmentation on both the predicted and the ground-truth IF images and then identify each prediction as one of the four possible detection outcomes, including the true positive (TP), true negative (TN), false positive (FP), and false negative (FN), from which we compute the cell-level detection metrics, including the sensitivity and specificity (see Materials and Methods). As summarized in Fig. 3C, the cell-level proliferation prediction achieves 83.55% sensitivity and 90.92% specificity; the apoptosis prediction achieves 75.91% sensitivity and 98.88% specificity. Notably, the scattering features in the proliferating cells cannot be easily perceived from the raw reflectance images, yet our DL model can capture the salient structural features with high accuracy. The balanced high sensitivity and specificity validates the reliability of our DL models for identifying these highly selective cell states/events.

Overall, these individual label prediction results validate our hypothesis that the improved sensitivity and resolution in reflectance images contain rich morphological features that can be utilized effectively for structural phenotyping by DL.

### Multiplexed prediction recovers biological accurate cellular structures

Next, we demonstrate the digital multiplexing capability by feeding the same reflectance input to each network and make six different IF predictions in parallel. By doing so, multiple subcellular structures and cell states are revealed simultaneously. In Fig. 4A, the image multiplexes the nucleus, Golgi apparatus, actin, and endosome virtual IF labels in a single wide field-of-view (FOV) for a large cell population. In Fig. 4B, the virtual labels for proliferating and apoptotic cells are multiplexed with the darkfield reflectance input in the same FOV as in Fig. 4A. These multiplexed predictions are performed on the cell batch under the Golgi apparatus staining condition. To further demonstrate the robustness of this digital multiplexing procedure, we show additional examples of multiplexed predictions performed on different cell batches/staining conditions in Supplementary Materials Fig. S6.

**Fig. 4.**
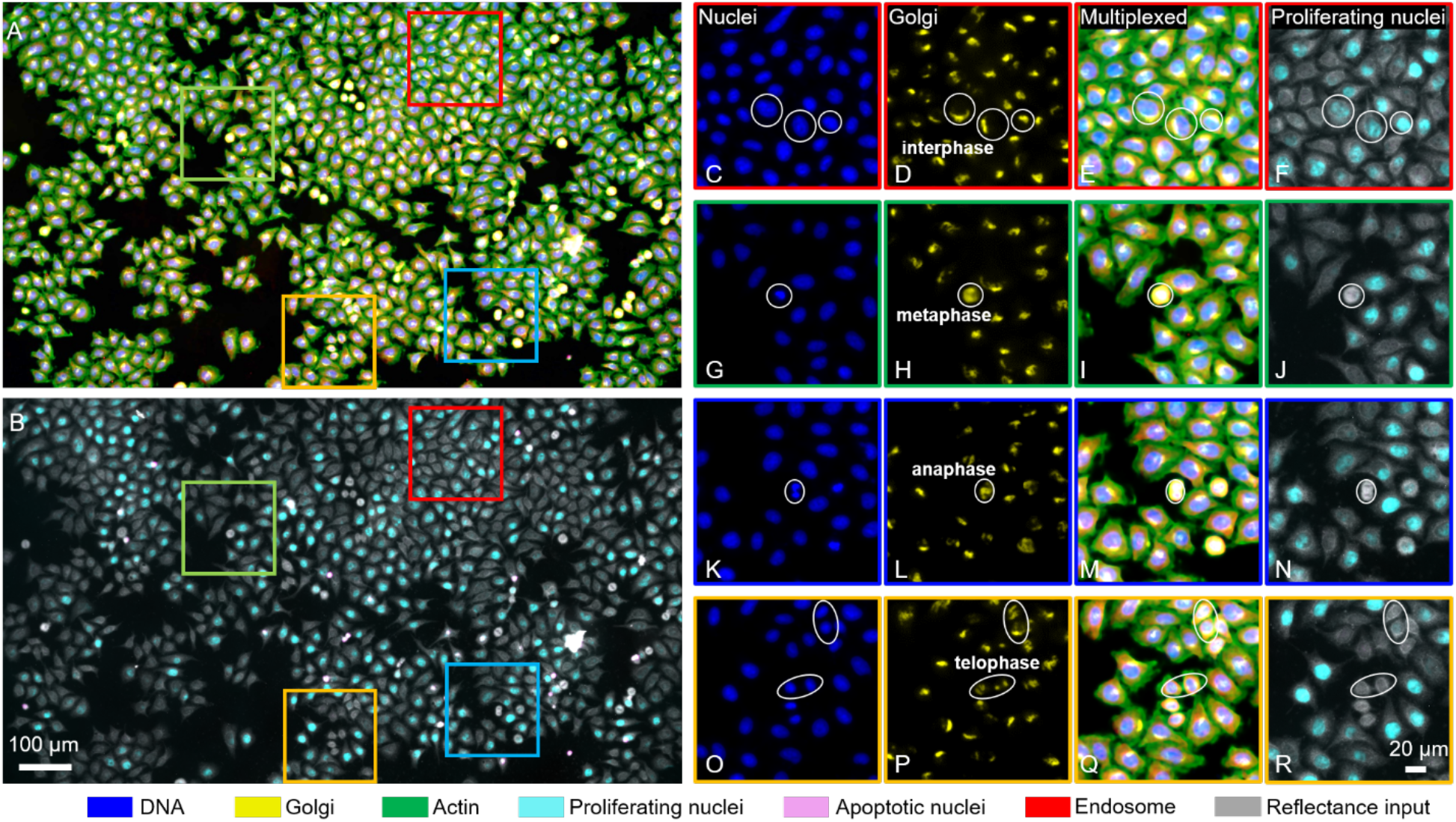
Multiplexed prediction on six IF labels from the same label-free input. **(A)** Visualization of the Full-FOV multiplexed prediction including DNA (blue), endosome (red), actin (green), and Golgi apparatus (yellow), and **(B)** proliferation (cyan) and apoptosis (magenta) from the same reflectance input (grayscale). **(C-R)** Zoomed-in of DNA, Golgi apparatus, multiplexed, and proliferation predictions. White circles indicate representative cell morphology during different phases of the cell cycle, including **(C-F)** interphase, **(G-J)** metaphase, **(K-N)** anaphase, and **(O-R)** telophase.

Importantly, our results show that the DL model can correctly capture and predict characteristic subcellular features during the cell cycle. During interphase, the nuclei have a regular rounded shape with nucleoli present, and Golgi apparatus is anchored primarily to one side of the nuclei (Figs. 4C-4E). At this stage, cells that have initiated or ongoing DNA/chromatin replications have a positive signal for proliferation (Fig. 4F). When cells enter mitosis, the chromosomes start to condense toward the centers of the cells, and nucleoli disappears. Golgi apparatus undergoes vesiculation and fragmentation, and its components are found scattered throughout the cytoplasm in the form of tiny (~50-nm) vesicles, often referred to as the “Golgi haze” (*25*). During metaphase, chromosomes align at the metaphase plate and the cell shape also changes dramatically, bulging into a sphere (Figs. 4G-4I). Golgi haze appears rounded, with a shaded center where chromosomes are located (Fig. 4H). During anaphase, the duplicated chromosomes separate from one another and move to opposite poles of the spindle (Fig. 4K-4M). During telophase, chromosomes start to de-condense, and begin to take on a more interphase-like shape (Figs. 4O-4Q). In this stage, Golgi apparatus has also completed replication, and reassembled into two closely-spaced cell bodies, referred to as “Golgi twins” (*25*) (Fig. 4P). There is no DNA replication during mitosis, so the markers for proliferation (incorporation of the fluorescent nucleoside, EdU) are absent for the metaphase, anaphase, and telophase (Figs. 4J, 4N, 4R). As shown in Fig. 4C-4R, these structural, subcellular, cell cycle-dependent features are accurately captured and predicted by our DL model, which validates our hypothesis that label-free reflectance imaging and DL enable structural phenotyping.

### Cell profile analysis on multiplexed images allows phenotyping and quantitative cytometry

A distinct advantage of multiplexing several markers is that it enables the development of multi-variant quantitative metrics for imaging cytometry and phenotyping across the large cell population and at the single-cell level. As a demonstration, we evaluate several cellular features, including cell morphology and fluorescence intensity, on the digitally multiplexed readouts. First, we generate fluorescence intensity scatter plots similar to those used in the flow cytometry, of EdU vs. Hoechst for all cells from the ground-truth and digitally multiplexed IF images in Fig. 5B and 5C, respectively. Evaluating the scatter plots on the population level, the DL-multiplexed prediction matches well with the ground truth, both of which show the increase in EdU and doubling of Hoechst intensity in the S and G2/M phases of the cell cycle, respectively. Next, we further evaluate the results on the individual cell level by overlaying the detection outcome for proliferation of every cell onto the same scatter plot in Fig. 5D. Specifically, by comparing the predicted proliferation IF label and the ground truth, each data point is labeled as one of the four detection outcomes (TP, TN, FP, or FN) (see details in Materials and Methods). Our analysis shows that the incorrect predictions (including FP and FN) tend to cluster around the boundaries between the S phase and S1 phases, leading to confusions in the DL predictions. There are relatively fewer incorrect predictions in G2/M phase, which are expected because of the distinctive morphological features during mitosis that can be easily captured by the DL model (*e.g.* metaphase, anaphase, and telophase in Fig. 4).

**Fig. 5.**
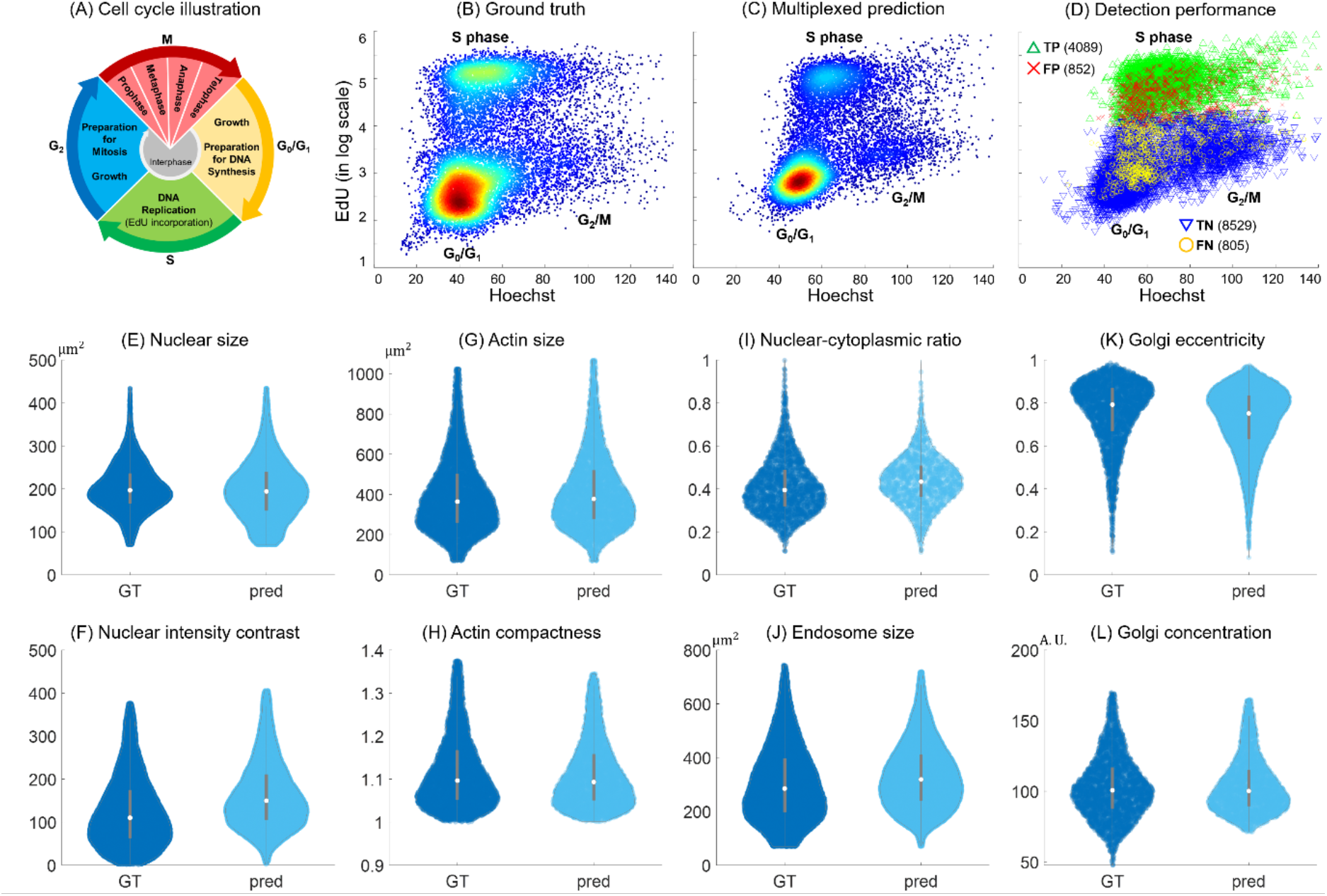
Cell profile analysis on digital multiplexed IF staining. **(A)** Illustration of the cell cycle. **(B-D)** The scatter plots for the whole cell-level EdU (proliferating DNA) and Hoechst (DNA) concentrations from **(B)** the co-stained ground-truth, **(C)** the DL-prediction, and **(D)** the detection performance for the EdU predictions quantified by TP, TN, FP, and FN, across the entire cell population under the proliferation staining condition. The total numbers of sample cells collected from the ground truth and DL-predictions are 14778 and 14275, respectively. The numbers of TP, FP, TN and FN in **(D)** are 4089, 852, 8529, and 805, respectively (in the brackets). **(E-L)** The comparisons of the statistics of eight single-cell profile metrics extracted from the entire cell population in the ground truth (GT) and the DL-predictions (pred), including **(E)** nuclear size, **(F)** DNA (nuclear) fluorescence intensity contrast, **(G)** cell (actin) size, **(H)** compactness of the actin, **(I)** NCR measured by the area ratio between the nuclei and actin, **(J)** endosome size, **(K)** eccentricity of the Golgi apparatus distribution, and **(L)** concentration of the Golgi apparatus. The total numbers of sample cells collected from the ground truth and DL-predictions are respectively 20021 and 25257 in **(E)** and **(F)**, 6183 and 5151 in **(G)** and **(H)**, 2246 and 1491 in **(I)**, 15944 and 22408 in **(J)**, 3380 and 9392 in **(K)**, 3380 and 3380 in **(L)**. All the single-cell profile metrics show good agreements between the predictions and the ground truths.

In Figs. 5E-5L, we extract several biologically relevant single-cell profile metrics using the predicted IF labels and compare them with the ground truths (see Materials and Methods), as visualized in the violin plots. In particular, we show the statistics of eight different morphological and subcellular structural parameters. In Fig. 5E-5F, we gather statistical data about the DNA label to measure the nuclear size and intensity contrast. In Fig. 5G-5H, we evaluate the actin size (i.e. cell size) and its compactness. In Fig. 5I, we compute the nuclear-cytoplasmic ratio (NCR), an important marker for cancers, as the area ratio between the nucleus and actin. In Fig. 5J, we measure the endosome size. In Fig. 5K-5L, we collect morphological parameters about the Golgi apparatus, including the eccentricity and concentration. Additional metrics are provided in the Supplementary Materials Fig. S7. For all these single-cell profile metrics, the prediction and the ground truth show excellent agreement. These results clearly demonstrate that our DL-augmented label-free cytometry can provide comprehensive morphological quantifications with high accuracy at the singlecell level, which is the key element for phenotyping and high-content screening (*26*).

### Saliency map reveals inner mechanism of the deep neural network

Deep neural networks have shown high expressivity for complex models, but suffer from poor explainability. Many theoretical explanations for the DL model have resorted to statistical perspective while treating the overall model as a “black box”. Instead, we utilize the “attention”-based technique (*24*) to elucidate on the specific label-free subcellular features that contribute to the fluorescence prediction. To do so, we treat the trained DL model as a mapping function between the input and the output. We then visualize the network’s gradient with respect to the input and extract the salient features (i.e. those having the largest gradients) the network pays most attention to (see details in Materials and Methods). The resulting “saliency map” highlights the most important features contributing to the IF prediction. By doing so, the saliency map directly evaluates the *specificity* of the structural features extracted from the reflectance images and how they are transformed to the target fluorescence labels by our network.

Figure 6 shows the computed saliency maps for each network across different sample batches and labeling conditions. Importantly, distinct subcellular features not only are highlighted by the network’s saliency map, but also have good visual correspondence to the targeted fluorescence label. By inspecting different columns, we show morphologically distinct features from different networks, indicating that different networks can indeed learn to recognize and focus on specific features present in the label-free images. For example, the DNA saliency maps show emphasis on nuclear boundaries and some subcellular structures. The actin saliency maps show concentration over the whole cell and spreading out to the membrane boundaries. The saliency maps for Golgi apparatus generally form shapes in “partial-moon” or circular lines. By contrast, the saliency maps for proliferation and apoptosis show that the network selectively pays attention to certain features around the nuclei. In addition, the saliency maps show that our network learns to extract *invariant* structural features specific to the underlying fluorescence label regardless of the cell preparation and labeling processes. Across different rows, we observe consistent saliency maps under different sample batches / staining conditions for the same labeling network.

**Fig. 6.**
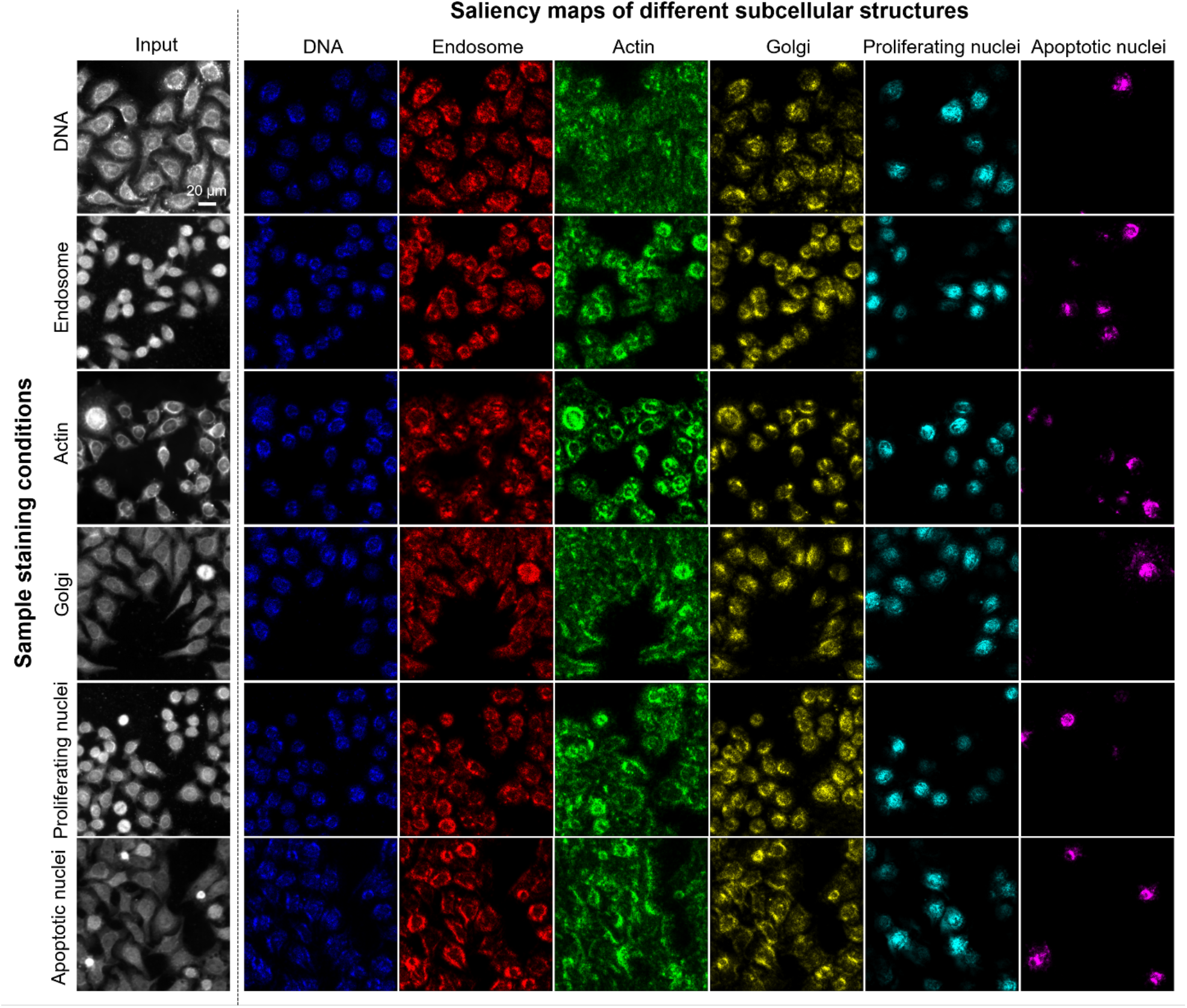
Saliency maps from each network across six cell batches with different staining conditions. The columns show the label-free input (the first darkfield reflectance channel) and the saliency maps for six different IF labels, including DNA (blue), endosome (red), actin (green), Golgi apparatus (yellow), proliferation (cyan), and apoptosis (magenta). The rows show the label-free input and the saliency maps from six cell batches under different staining conditions. The saliency maps show good consistency across different batches and highlight distinct morphological features.

## Discussion

In this study, we have presented a DL-augmented label-free cytometry technique that accurately predicted six fluorescence targets in-parallel at the single-cell level. The accuracy has been improved up to 3× in predicting subcellular structures as compared to the current state-of-the-art. Remarkably, the DL model is able to accurately recognize subcellular organelles, such as Golgi apparatus reconfigure during the cycle of proliferation, as well as to distinguish subtle morphological differences between the proliferating and nonproliferating cells. These results demonstrate the data-driven model’s unique capability of holistically extracting “non-intuitive” structural features from the label-free imaging data on a large cell population. The *specificity* for predicting cellular features by our DL model is illuminated by the saliency maps. This analysis demonstrated the ability of the DL models in processing highly complex and entangled structural information from scattering images.

Beyond predicting IF labels, we have further demonstrated quantitative cytometry analysis based on the multiplexed digital output from our DL models. Importantly, our analysis has shown that a multitude of single-cell profile metrics can be accurately extracted from the DL predictions. The digital multiplexing enabled us to simultaneously quantify several morphological features on multiple subcellular components across a large cell population. This capability drastically improves the technique’s throughput for structural phenotyping in the application of imaging cytometry, such as high-content analysis/screening. Cell morphological features are effective phenotypes for different disease states and environmental influences. This phenomenon is well described and practiced in pathology and cell biology. Nuclear condensation, enlargement, and increased NRC are ubiquitous hallmarks of cancers (*5, 6*). Cell morphology is distinct for different cell types, which is often denoted in their terminology (i.e. astrocytes, macrophage, squamous, and columnar cells, *etc.*), and stem cells change structures along separate differentiation paths (*13*). It has been shown that cell morphological changes can be directly associated with changes of morphogenic gene expressions (*27*), and comprehensive morphological profiling can be used to detect genetic functions (*28*). Meanwhile, it has been shown that DL techniques can holistically capture complex structural features for classification. This has found broad applications in detecting cell types (*13, 14, 29*), cell states (*15–18, 22*), drug response (*19*), and stem cell lineage (*20*). By fully leveraging the label-free and high multiplexing nature of our technique, it can potentially generate significant impacts in imaging cytometry by offering unprecedented information content and discovering new compound morphological features necessitating multiplexed fluorescence readouts.

Label-free, DL-augmented method of cell-morphology profiling is data-driven and ultimately relies on the rich information content in the images. Our LED-array reflectance microscopy enables multi-contrast imaging (*i.e.* angle-dependent darkfield and drDPC) by detecting the angled-dependent backscattering signals by a programmable LED array without any mechanical moving parts. The superior sensitivity in detecting subtle structures using backscattering than transmission-microscopy is well documented (*2*). The superb sensitivity of backscattering-based method has been demonstrated in a variety of techniques, such as partial wave spectroscopy (*30*), confocal light absorption and scattering spectroscopic microscopy (*31*), confocal reflectance quantitative phase microscopy (*32*), and spatial-domain low-coherence quantitative phase microscopy (*33*). In addition, the angle-dependent measurement has been used to measure characteristic structural length scale (*6*) and to enable 3D reconstruction of the refractive index distribution (*34*). Leveraging angle-dependent reflectance signal, we outperformed the state-of-the-art for predicting multiple subcellular components. By switching to a higher NA objective lens, the prediction performance of our DL model to subcellular structures, such as those missed in our actin label predictions, may be further improved. Recently, the LED array microscopy has also been extensively explored in transmission that allows sampling the low spatial frequency components in the Fourier space (*34*). In addition, backscattering spectroscopic techniques further enable characterization of ultrastructural phenotypes with sensitivity down to nm-length scale (*35*). A potential future improvement of our imaging system is to incorporate additional transmission and multispectral LED-array illumination to fully exploit the angle- and wavelength-dependent scattering contrast with a single objective lens by versatile illumination engineering.

One limitation of our current work is that it is based on fixed cells that does not allow longitudinal imaging. This can be overcome by using fluorescent reporter cell lines or live cell dyes to provide the fluorescence ground-truth (*10*) and enable dynamic observation. The additional temporal dimension may further improve the model’s sensitivity in cell phenotyping and discover new label-free features by incorporating the information about the cell dynamics (*20, 22, 36*). Another limitation of the DL framework we used here is that it cannot be generalized to different types of cells. Techniques based on transfer learning (*37, 38*) and domain adaptation (*39*) will be investigated in our future work to overcome this limitation.

The variations in the prediction accuracy of our DL models are currently evaluated *post hoc*, and based on pixel-wise and cell-level metrics by comparing the DL predictions and the ground truth. For many biomedical applications, it is beneficial to understand how much error the model may make without knowing the ground truth, *i.e.* the confidence of the model predictions. Emerging Bayesian DL based uncertainty quantification techniques have proved useful to provide a proxy estimate of the prediction accuracy and quantify the model confidence (*40, 41*), which will be adapted in our future work.

In summary, we have reported a label-free imaging cytometry technique that multiplexes six IF labels in-parallel with high accuracy via DL models. We have validated the fluorescence predictions by comparing them to the ground-truth IF images. In addition, we have conducted imaging cytometry studies on several quantitative morphological metrics on subcellular structures and phenotyping of cell proliferation. Finally, the specificity of the DL model is assessed by visualizing the saliency map at the single cell level across different staining and fixation conditions. With this unique combination of new capabilities, this new framework may find wide applications in image-based cytometry, in particular for high-content screening and analysis.

## Materials and Methods

### Cell preparation and immunofluorescence staining

HeLa cells were cultured in a Dulbecco’s Modified Eagle Medium (DMEM) with 10% fetal bovine serum (Gibco, 10564011) and 5% penicillin streptomycin (Gibco, 15140122). The cells were trypsinized and passaged twice a week. Two days before the staining and imaging, cells were cultured on glass-bottom petri dishes (FluoroDish FD35-100) which were first treated with 10% poly-L-lysine (SigmaAldrich, RNBG0769) with PBS for ten mins in an incubator. The staining and imaging were performed on the glass-bottom dishes.

We follow the standard IF staining protocols. In total, six IF stains are used to label DNA (Hoechst), actin (Phalloidin, Alexa Fluor 488 Phalloidin, Invitrogen, A12379), endosome (EEA1, Santa Cruz, sc-137130 AFF488), Golgi apparatus (GM130, Cell Signaling, 12480), proliferation (EdU, Click-iT Plus EdU Alexa Fluor 488 kit, Invitrogen), and apoptosis (TUNEL, Click-iT TUNEL Alexa Fluor 488 kit, Invitrogen). The HeLa cells were first fixed with ice-cold methanol, washed three times (10 min each) in 0.05% PBST (0.05% Triton X-100 PBS solution), and incubated for 20 mins at room temperature in a blocking solution containing 0.25% Triton X-100 and 10% bovine serum albumin in PBS. Alexa-488 conjugated antibodies were diluted in the blocking solution with the recommended concentration by the manufacturers and incubated with cells to label the specific subcellular components (EEA1 for endosome, Phalloidin for actin). For actin staining, cells were fixed with ice-cold acetone to preserve the structures. For Golgi staining, a secondary antibody Anti-rabbit IgG (Santa Cruz, 4412S) was diluted in blocking solution and used to culture the cells for 1.5 hours at room temperature in dark. To stain cell proliferation and apoptosis, we used EdU and TUNEL assays, respectively, according to the recommended protocol by Invitrogen. The apoptosis was induced by culturing the cells with 1μM Staurosporine for 24 hours. In all the above stains, cell nuclei were counterstained with 1× Hoechst 33342.

### Image data acquisition

We collect the data using our custom-built multimodal reflectance microscope (*8*), as shown in Fig. 1A. A custom-built LED array consisting of two LED rings is used for providing controllable darkfield illumination in reflection. We use commercially available LEDs (APTF1616SEEZGQBDC, Kingbright) that can provide three independent RGB color channels (central wavelength is 460, 515, and 630 nm, respectively). All the LEDs are individually addressable using two cascaded LED drivers (TLC5955, Texas Instruments). A microcontroller (Teensy 3.2, PJRC) provides the camera trigger signal through digital Input/Output pins and simultaneously controls the LED illumination pattern. The LED array is mounted around the objective lens (10× 0.3 NA, UPlanFL N, Olympus, Japan) using a 3D-printed adapter. The tube lens is a commercial SLR lens (Nikon AF DC-NIKKOR 135 mm f/2D) to maximize the FOV. The microscope provides an overall 7.5× magnification. An sCMOS camera (CS2100M-USB, Thorlabs, 1920 × 1080 pixels, 5.04-μm pixel size, 16-bit depth) is used to acquire the images. We capture four darkfield reflectance images by using half-annulus green LED patterns along different orientations, including top (*I*_Top_), bottom (*I*_BOttOm_), left (*I*_Left_), and right (*I*_Right_). The exposure time is 700 ms. Two drDPC images along two orthogonal orientations are generated by

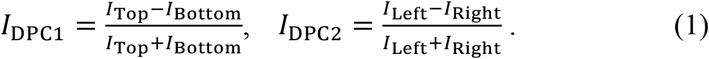

The fluorescence excitations are provided by two LED sources (M365LP1 and M470L4, Thorlabs, central wavelength 365 nm, 470 nm, respectively) combined with a dichroic mirror (DMLP425R, Thorlabs). The epi-fluorescence illumination is introduced by a 50/50 beam splitter (CCM1-BS013, Thorlabs). The emission filters (MF460-60, MF525-39, Thorlabs) are placed on a filter wheel (CFW6, Thorlabs) for blue and green fluorescence emissions. Two-channel fluorescence images are acquired sequentially after acquiring the reflectance images. The exposure time is 400 ms for IF imaging. Specifically, the first green channel is for one of the five IF antibodies conjugated with the green fluorophores (Alexa 488) for endosome, actin, Golgi apparatus, proliferation, and apoptosis; the second blue channel is for the co-stained DNA. We capture 30 image stacks for each sample batch / IF stain.

### Data preprocessing procedure

The raw reflectance and fluorescence images are preprocessed before feeding into our deep neural networks for training. The preprocessing procedure consists of four steps, including flat-field cropping, image denoising, background correction, and intensity normalization. Since the fluorescence excitation illumination is not evenly distributed across the entire rectangular FOV, we first perform flat-field cropping by using only the central 1000 × 1080-pixel region for training, where the excitation is approximately uniform. Second, we perform image denoising on the measurements. We apply two denoising approaches. In the first approach, we apply an unsupervised DL-based denoising algorithm, noise2void (*42*), to suppress the sensor noise present in the images. To do so, each 1000 × 1080-pixel image is cropped into 256 × 256-pixel patches. Each image patch is fed to a blind-spot network to perform denoising. After denoising, the patches are then stitched back together by alpha blending. This unsupervised denoising algorithm is found to be effective in removing unstructured, signal-independent noise, including the sensor noise and isolated hot pixels, in particular for measurements with low signal-to-noise ratios (SNRs). The whole training and inference (denoising) procedure takes ~10 hours for processing the entire dataset containing 30 images. We find this denoising procedure is only necessary for processing the Golgi and proliferation fluorescence images, as well as for the reflectance images for the actin prediction where the sensor noise severely corrupts the images. In the second approach, when the SNRs are sufficiently high for the measurements on other cell batches, we use a computationally more efficient morphology opening operation to remove the hot pixels in the fluorescence images under the assumption that hot pixels are isolated pixels with extreme intensity values. The opening operation takes a square kernel of size 2 × 2 pixels. This hot-pixel removal procedure takes ~15 min to process the entire dataset containing 30 images. Third, we perform background correction on the fluorescence images by eliminating the potential background bias across the batches. To do so, we calculate the histogram of each fluorescence image and denote the mode value (i.e. the most frequent value) as the constant background. This background of each fluorescence image is subtracted; the negative values from the subtraction are clipped to zero. Fourth, we perform intensity normalization by normalizing the pixel values of both the input and output images to be between 0 and 1. Additional details about the data preprocessing steps are shown in Supplementary Materials Fig. S2.

Our network takes 256 × 256-pixel input images. Accordingly, we split the 30 pairs of images into 256 × 256-pixel patches to generate the training and testing data. For each IF prediction, 512 training samples and 128 testing samples are randomly generated from the full-FOV image pairs. Each input stack consists of four different channels combining darkfield reflectance images from different oblique illumination patterns (Fig. 1A). In addition, we construct the two-direction drDPC images using Equation (1) from the darkfield images (Fig. 1A), which were found particularly effective for predicting the actin, proliferation, and apoptosis labels. We provide example comparisons of the prediction results with and without the drDPC inputs for all six IF labels in Supplementary Materials Fig. S3. Quantitative comparisons of the prediction results with and without the drDPC inputs for all six IF labels are provided in Supplementary Materials Table S1. When making the full-FOV predictions (e.g. Fig. 4), we use the entire 1080 × 1920-pixel reflectance images since they do not suffer from the non-uniform illumination issue.

### Neural network implementation

We develop a convolutional neural network (CNN) to learn the highly complex nonlinear mapping between the morphology information contained in the multi-channel reflectance images and the fluorescence labels. The network structure follows the encoder-decoder “U-net” architecture and further incorporates the dense-blocks and skip-connections to enable high-resolution information prediction (*40*). The input of the preprocessed 256 × 256-pixel reflectance image stack passes through the “encoder” path consisting of four dense blocks followed by the max-pooling layers, and the bottleneck feature maps are then fed into the “decoder” path with four dense blocks followed by upsampling layers. The skip connections bridge the lower-level activation maps with higher-level activation maps and preserve the high-frequency information. More details about the network are provided in Supplementary Materials Fig. S1. We use the negative PCC (NPCC) as the training loss (*43*). We train our network using ADAM optimizer with 500 epochs and 0.1 training/validation splitting. No overfitting is observed during the training.

### Quantitative evaluation of network prediction

We use the PCC to evaluate the performance of the regression-type of problems. Specifically, the PCC is used to quantify the prediction quality for pervasive subcellular features, including DNA, endosome, actin, and Golgi apparatus labels. It computes the statistical correlation between the predicted and ground-truth IF image patches and is able to quantify the pixel-level similarity on the fine subcellular features. The PCC between the prediction *X* and the ground truth *Y* (each image is reshaped to a *N* dimensional vector) is computed as

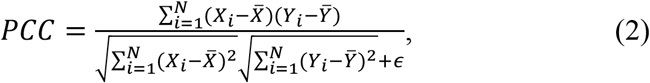

where *ϵ* = 10^-10^ is a small regularizer to prevent zero denominator, 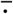 denotes the mean, and *i* is the index of each vector. The value of the PCC ranges from –1 to +1, where ±1 indicates total positive or negative correlation and 0 indicates no correlation. The PCC computation is implemented by a custom code in Python. The PCCs are computed on the testing image patches, each containing a 128×128-pixel FOV. In total, the statistics from 676 patches from four large FOV testing images of size 908×908 pixels at each staining condition are aggregated and shown in the violin plots in Fig. 3B.

To evaluate the spatial variations of the prediction performance, we construct the PCC map for each label prediction. To do so, 169 consecutive image patches are obtained from each large FOV image by cropping the image with a 128×128-pixel sliding-window and 64×64-pixel overlap between the neighboring patches. The patch-wise PCC map is computed and shown in Supplementary Materials Fig. S5. The lower PCC patches are found to generally align with the background region where very low reliable IF readings are present. We treat these patches as outliers. Accordingly, we use a median-based outlier detection and removal algorithm (see Supplementary Material Section S1) to remove these background outliers when constructing the violin plots in Fig. 3.

We use the AUC to quantify the detection performance for identifying the selective cell events including the proliferation and apoptosis. To plot the ROC of the detection label performance, the ground-truth labels are first binarized with certain thresholds. We use the well-established Otsu’s method to reliably compute the binarization threshold values (*44*), which finds the optimal values that maximize the intensity variance between the signals and the background based on a histogram clustering criterion. Specifically, the threshold values are calculated based on the aggregated histogram from the entire ground-truth dataset of proliferation and apoptosis, and are found to be 0.31 and 0.11, respectively. Next, each continuous-valued pixel in the predicted image is regarded as the predicted probability of expressing the IF label at this pixel. By using the binarized ground-truth images as the target, the predictor (the trained CNN) achieves different pixel-wise True Positive Rate (TPR) and False Positive Rate (FPR) under different detection thresholds on the predicted images. By varying the detection thresholds, the TPR and FPR as functions of the thresholds can be plotted on the ROC curve. The AUC measures the area under the ROC curve and provides an aggregated quantification of the performance across all possible detection thresholds. The AUC is computed by the built-in functions ‘roc_curve’ and ‘auc’ in the scikit-learn module in Python. The AUCs are computed on the testing image patches, each contains a 128 × 128-pixel FOV. In total, the statistics from 676 testing image patches from four large FOV testing images of size 908×908 pixels at each condition are aggregated and shown in the violin plots in Fig. 3B.

We also quantify the single-cell level detection accuracy for the proliferation and apoptosis label predictions. To do so, we develop an automatic image processing pipeline to segment and identify each prediction as TP, TN, FP, or FN in CellProfiler. Subsequently, we compute the cell-level detection metrics, including the sensitivity and specificity. The sensitivity (a.k.a recall) is computed from the TPR, and the specificity (a.k.a selectivity) is computed as the TNR as

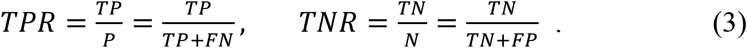

The implementation details of the image processing pipeline are described in ‘Digital cytometry analysis’ section and Supplementary Material Section S2 and Fig. S8.

### Digital cytometry analysis

We develop a digital cytometry analysis framework for exploring the interdependencies of different fluorescence markers on the multiplexed predictions. A commonly used flow cytometry analysis is performed by displaying the scatter plot of single-cell level proliferating DNA concentration in the log-scale against the DNA concentration in the linear-scale. Different from flow cytometry that directly collects the integrated fluorescence intensity from each cell, our method performs imaging with subcellular resolution across a large cell population. As a result, we first perform cell segmentation and then aggregation of the fluorescence signals within each cell region to carry out the single-cell digital cytometry analysis. We perform the digital cytometry analysis to relate the cell-level Hoechst and EdU fluorescence concentration using CellProfiler (*45*). Since our data contain co-registered two-channel fluorescence images with co-stained Hoechst and EdU, we can directly compare the ground-truth cytometry scatter plot with that from our multiplexed prediction. To generate the scatter plots, we process the co-registered Hoechst and EdU images and the multiplexed predictions using CellProfiler with a standard single-cell segmentation-based pipeline and extract the paired cell-level fluorescence intensity data. A small value (10^-10^) is added to the fluorescence intensity of the proliferating DNA before taking the log operation to avoid the singularity at 0. Three distinct clusters representing the S, G1, and G2/M phase are clearly shown in the digital cytometry scatter plots. Additional details are provided in Supplementary Material Section S3 and Fig. S9.

We construct an image processing pipeline to quantify the single-cell-level detection performance on the proliferation and apoptosis labels (denoted as the target IF in this paragraph). First, for each of the cell event labels, we perform segmentation on the costained DNA channel ground truth images to find all the nuclear localizations. Second, we segment all the nuclei in both the target IF prediction and ground-truth images. Third, each nucleus is labeled as one of the four possible detection outcomes. Specifically, a prediction is TP if both the prediction and the ground truth express the target IF. A prediction is TN if neither the prediction nor the ground truth express the target. A prediction is FP if the prediction expresses the target IF while the ground truth does not. A prediction is FN if the prediction does not express the target IF while the ground truth does. To further investigate the effect of false detections (FP/FN) on the prediction digital cytometry scatter plot, we apply the above pipeline and label each prediction as TP, TN, FP, or FN in Fig. 5D. Additional details are provided in Supplementary Material Section S2.

### Cell profile analysis

We use CellProfiler (*45*) to generate the single-cell profiles across each fluorescence image. We feed the ground truth and the predicted IF images of DNA, actin, endosome, and Golgi apparatus to CellProfiler. After initial cell segmentation, single-cell level parameters of morphology and intensity distribution are computed automatically by different measurement modules in CellProfiler, including the fluorescence marker size (area), compactness, eccentricity, fluorescence concentration, and single-cell-level fluorescence variance and contrast. In addition, we compute the compound metric, NCR, based on the multiplexed DNA and actin fluorescence labels. To compute the NCR, co-registered ground-truth / prediction images containing the DNA and actin labels are processed individually in CellProfiler. The NCR is computed as the ratio between the area of segmented nuclei and actin masks. The single-cell profiles also present the same outliers from the background regions, which are eliminated by our outlier removal algorithm (see Supplementary Material Section S1) when constructing the violin plots. Additional details are provided in Supplementary Material Section S4.

### Saliency map visualization

Our network performs image-to-image translation and can be treated as a function relating the input and output images. On the other hand, the norm of the network output is a scalar function. As a result, computing and visualizing the gradient over the norm is still possible. Based on this notion, we compute the saliency map as the gradient of the norm of the output with respect to the given input image. The practical implementation is achieved by the automatic differentiation feature in TensorFlow, as detailed in Supplementary Materials Section S5. We specified the gradient modifier as the absolute values of the gradient, which shows the regions in the input that contribute most to the change in the output regardless of the sign of the change (i.e. negative or positive). We also used the guided backpropagation to propagate only the positive gradients for positive activations to achieve a smoother visualization. The saliency map was computed for each network and on different sample input (for varying sample batches / fixation conditions). The inputs are 256 × 256-pixel image stacks randomly selected from the testing groups under the six sample conditions. The computed saliency maps are normalized to have a uniform range between 0 and 1 for visualization.

## Acknowledgments

We thank Alex Matlock for helpful discussions on cell phenotyping visualization and evaluation, and Boston University Shared Computing Cluster for proving the computational resources.

## Funding

This work was supported by the National Science Foundation (1846784), National Institutes of Health (R01NS108464), and Boston University Hariri Institute Research Incubation Award.

## Author contributions

L.T. and J.Y. conceived the idea. S.F. and Y.K. prepared the cell sample culturing, fixation, staining and acquired all the imaging data. S.C., Y. L. and Y. X. conducted the image processing, network training, cell phenotype and profile analysis, quantitative evaluation and saliency map analysis. W.S. developed the LED-array reflectance microscope platform. L.T., J.Y. and S.C. further discussed the results and refined the deep learning model, cell profiling and cytometry pipeline. All authors contributed to the writing of the manuscript.

## Competing interests

The authors declare that they have no competing interests.

## Data and materials availability

All data needed to evaluate the conclusions in the paper are present in the paper and/or the Supplementary Materials. The data that support the findings in this study are available upon request from the corresponding authors.

## Notes

### Competing Interest Statement

The authors have declared no competing interest.

## References and Notes

1. F. Zanella, J. B. Lorens, W. Link, High content screening: seeing is believing. Trends Biotechnol. 28, 237–245 (2010).

2. N. N. Boustany, S. A. Boppart, V. Backman, Microscopic Imaging and Spectroscopy with Scattered Light. Annu. Rev. Biomed. Eng. 12, 285–314 (2010).

3. R. Kasprowicz, R. Suman, P. O’Toole, Characterising live cell behaviour: Traditional label-free and quantitative phase imaging approaches. Int. J. Biochem. Cell Biol. 84, 89–95 (2017).

4. Y. Park, C. Depeursinge, G. Popescu, Quantitative phase imaging in biomedicine. Nat. Photonics. 12, 578 (2018).

5. S. Uttam, H. V. Pham, J. LaFace, B. Leibowitz, J. Yu, R. E. Brand, D. J. Hartman, Y. Liu, Early Prediction of Cancer Progression by Depth-Resolved Nanoscale Mapping of Nuclear Architecture from Unstained Tissue Specimens. Cancer Res. 75, 4718–4727 (2015).

6. A. Wax, K. J. Chalut, Nuclear Morphology Measurements with Angle-resolved Low Coherence Interferometry for Application to Cell Biology and Early Cancer Detection. Anal. Cell. Pathol. Amst. 34, 207 (2011).

7. L. Cherkezyan, I. Capoglu, H. Subramanian, J. D. Rogers, D. Damania, A. Taflove, V. Backman, Interferometric spectroscopy of scattered light can quantify the statistics of subdiffractional refractive-index fluctuations. Phys. Rev. Lett. 111, 033903 (2013).

8. W. Song, A. Matlock, S. Fu, X. Qin, H. Feng, C. V. Gabel, L. Tian, J. Yi, LED array reflectance microscopy for scattering-based multi-contrast imaging. Opt. Lett. 45, 1647–1650 (2020).

9. A. Matlock, A. Sentenac, P. C. Chaumet, J. Yi, L. Tian, Inverse scattering for reflection intensity phase microscopy. Biomed. Opt. Express. 11, 911–926 (2020).

10. C. Ounkomol, S. Seshamani, M. M. Maleckar, F. Collman, G. R. Johnson, Label-free prediction of three-dimensional fluorescence images from transmitted-light microscopy. Nat. Methods. 15, 917–920 (2018).

11. E. M. Christiansen, S. J. Yang, D. M. Ando, A. Javaherian, G. Skibinski, S. Lipnick, E. Mount, A. O’Neil, K. Shah, A. K. Lee, P. Goyal, W. Fedus, R. Poplin, A. Esteva, M. Berndl, L. L. Rubin, P. Nelson, S. Finkbeiner, In Silico Labeling: Predicting Fluorescent Labels in Unlabeled Images. Cell. 173, 792–803.e19 (2018).

12. S.-M. Guo, L.-H. Yeh, J. Folkesson, I. E. Ivanov, A. P. Krishnan, M. G. Keefe, E. Hashemi, D. Shin, B. B. Chhun, N. H. Cho, M. D. Leonetti, M. H. Han, T. J. Nowakowski, S. B. Mehta, Revealing architectural order with quantitative label-free imaging and deep learning. eLife. 9 (2020), doi:10.7554/eLife.55502.

13. D. Kusumoto, S. Yuasa, The application of convolutional neural network to stem cell biology. Inflamm. Regen. 39, 14 (2019).

14. D. Kusumoto, M. Lachmann, T. Kunihiro, S. Yuasa, Y. Kishino, M. Kimura, T. Katsuki, S. Itoh, T. Seki, K. Fukuda, Automated Deep Learning-Based System to Identify Endothelial Cells Derived from Induced Pluripotent Stem Cells. Stem Cell Rep. 10, 1687–1695 (2018).

15. C. L. Chen, A. Mahjoubfar, L.-C. Tai, I. K. Blaby, A. Huang, K. R. Niazi, B. Jalali, Deep Learning in Label-free Cell Classification. Sci. Rep. 6, 21471 (2016).

16. T. Blasi, H. Hennig, H. D. Summers, F. J. Theis, J. Cerveira, J. O. Patterson, D. Davies, A. Filby, A. E. Carpenter, P. Rees, Label-free cell cycle analysis for high-throughput imaging flow cytometry. Nat. Commun. 7, 10256 (2016).

17. P. Eulenberg, N. Köhler, T. Blasi, A. Filby, A. E. Carpenter, P. Rees, F. J. Theis, F. A. Wolf, Reconstructing cell cycle and disease progression using deep learning. Nat. Commun. 8, 463 (2017).

18. J. K. Zhang, Y. R. He, N. Sobh, G. Popescu, Label-free colorectal cancer screening using deep learning and spatial light interference microscopy (SLIM). APL Photonics. 5, 040805 (2020).

19. H. Kobayashi, C. Lei, Y. Wu, A. Mao, Y. Jiang, B. Guo, Y. Ozeki, K. Goda, Label-free detection of cellular drug responses by high-throughput bright-field imaging and machine learning. Sci. Rep. 7, 12454 (2017).

20. F. Buggenthin, F. Buettner, P. S. Hoppe, M. Endele, M. Kroiss, M. Strasser, M. Schwarzfischer, D. Loeffler, K. D. Kokkaliaris, O. Hilsenbeck, T. Schroeder, F. J. Theis, C. Marr, Prospective identification of hematopoietic lineage choice by deep learning. Nat. Methods. 14, 403–406 (2017).

21. Y. Jo, S. Park, J. Jung, J. Yoon, H. Joo, M. Kim, S.-J. Kang, M. C. Choi, S. Y. Lee, Y. Park, Holographic deep learning for rapid optical screening of anthrax spores. Sci. Adv. 3, e1700606 (2017).

22. A. Zaritsky, A. R. Jamieson, E. S. Welf, A. Nevarez, J. Cillay, U. Eskiocak, B. L. Cantarel, G. Danuser, bioRxiv, in press, doi:10.1101/2020.05.15.096628.

23. T. Ching, D. S. Himmelstein, B. K. Beaulieu-Jones, A. A. Kalinin, B. T. Do, G. P. Way, E. Ferrero, P.-M. Agapow, M. Zietz, M. M. Hoffman, W. Xie, G. L. Rosen, B. J. Lengerich, J. Israeli, J. Lanchantin, S. Woloszynek, A. E. Carpenter, A. Shrikumar, J. Xu, E. M. Cofer, C. A. Lavender, S. C. Turaga, A. M. Alexandari, Z. Lu, D. J. Harris, D. DeCaprio, Y. Qi, A. Kundaje, Y. Peng, L. K. Wiley, M. H. S. Segler, S. M. Boca, S. J. Swamidass, A. Huang, A. Gitter, C. S. Greene, Opportunities and obstacles for deep learning in biology and medicine. J. R. Soc. Interface. 15 (2018), doi:10.1098/rsif.2017.0387.

24. K. Simonyan, A. Vedaldi, A. Zisserman, Deep Inside Convolutional Networks: Visualising Image Classification Models and Saliency Maps. ArXiv13126034 Cs (2014) (available at http://arxiv.org/abs/1312.6034).

25. G. M. Gaietta, B. N. G. Giepmans, T. J. Deerinck, W. B. Smith, L. Ngan, J. Llopis, S. R. Adams, R. Y. Tsien, M. H. Ellisman, Golgi twins in late mitosis revealed by genetically encoded tags for live cell imaging and correlated electron microscopy. Proc. Natl. Acad. Sci. 103, 17777–17782 (2006).

26. J. C. Caicedo, S. Cooper, F. Heigwer, S. Warchal, P. Qiu, C. Molnar, A. S. Vasilevich, J. D. Barry, H. S. Bansal, O. Kraus, M. Wawer, L. Paavolainen, M. D. Herrmann, M. Rohban, J. Hung, H. Hennig, J. Concannon, I. Smith, P. A. Clemons, S. Singh, P. Rees, P. Horvath, R. G. Linington, A. E. Carpenter, Data-analysis strategies for image-based cell profiling. Nat. Methods. 14, 849–863 (2017).

27. C. Bakal, J. Aach, G. Church, N. Perrimon, Quantitative Morphological Signatures Define Local Signaling Networks Regulating Cell Morphology. Science. 316, 1753–1756 (2007).

28. M. H. Rohban, S. Singh, X. Wu, J. B. Berthet, M.-A. Bray, Y. Shrestha, X. Varelas, J. S. Boehm, A. E. Carpenter, Systematic morphological profiling of human gene and allele function via Cell Painting. eLife. 6, e24060 (2017).

29. S.-M. Guo, A. P. Krishnan, J. Folkesson, I. Ivanov, B. Chhun, N. Cho, M. Leonetti, S. B. Mehta, Revealing architectural order with polarized light imaging and deep neural networks. bioRxiv, 631101 (2019).

30. D. Damania, H. K. Roy, D. Kunte, J. A. Hurteau, H. Subramanian, L. Cherkezyan, N. Krosnjar, M. Shah, V. Backman, Insights into the field carcinogenesis of ovarian cancer based on the nanocytology of endocervical and endometrial epithelial cells. Int. J. Cancer. 133, 1143–1152 (2013).

31. I. Itzkan, L. Qiu, H. Fang, M. M. Zaman, E. Vitkin, I. C. Ghiran, S. Salahuddin, M. Modell, C. Andersson, L. M. Kimerer, P. B. Cipolloni, K.-H. Lim, S. D. Freedman, I. Bigio, B. P. Sachs, E. B. Hanlon, L. T. Perelman, Confocal light absorption and scattering spectroscopic microscopy monitors organelles in live cells with no exogenous labels. Proc. Natl. Acad. Sci. 104, 17255–17260 (2007).

32. V. R. Singh, Y. A. Yang, H. Yu, R. D. Kamm, Z. Yaqoob, P. T. C. So, Studying nucleic envelope and plasma membrane mechanics of eukaryotic cells using confocal reflectance interferometric microscopy. Nat. Commun. 10, 3652 (2019).

33. P. Wang, R. K. Bista, W. E. Khalbuss, W. Qiu, S. Uttam, K. D. Staton, L. Zhang, T. A. Brentnall, R. E. Brand, Y. Liu, Nanoscale nuclear architecture for cancer diagnosis beyond pathology via spatial-domain low-coherence quantitative phase microscopy. J. Biomed. Opt. 15, 066028 (2010).

34. R. Ling, W. Tahir, H.-Y. Lin, H. Lee, L. Tian, High-throughput intensity diffraction tomography with a computational microscope. Biomed. Opt. Express. 9, 2130–2141 (2018).

35. J. Yi, A. J. Radosevich, J. D. Rogers, S. C. P. Norris, İ. R. Çapoğlu, A. Taflove, V. Backman, Can OCT be sensitive to nanoscale structural alterations in biological tissue? Opt. Express. 21, 9043–9059 (2013).

36. Z. Wu, B. B. Chhun, G. Schmunk, C. N. Kim, L.-H. Yeh, T. Nowakowski, J. Zou, S. B. Mehta, bioRxiv, in press, doi:10.1101/2020.07.20.213074.

37. Y. Rivenson, H. Wang, Z. Wei, K. de Haan, Y. Zhang, Y. Wu, H. Günaydin, J. E. Zuckerman, T. Chong, A. E. Sisk, L. M. Westbrook, W. D. Wallace, A. Ozcan, Virtual histological staining of unlabelled tissue-autofluorescence images via deep learning. Nat. Biomed. Eng. 3, 466–477 (2019).

38. Y. Rivenson, K. de Haan, W. D. Wallace, A. Ozcan, Emerging Advances to Transform Histopathology Using Virtual Staining. BME Front. 2020 (2020), doi:https://doi.org/10.34133/2020/9647163.

39. R. Hollandi, A. Szkalisity, T. Toth, E. Tasnadi, C. Molnar, B. Mathe, I. Grexa, J. Molnar, A. Balind, M. Gorbe, M. Kovacs, E. Migh, A. Goodman, T. Balassa, K. Koos, W. Wang, J. C. Caicedo, N. Bara, F. Kovacs, L. Paavolainen, T. Danka, A. Kriston, A. E. Carpenter, K. Smith, P. Horvath, nucleAIzer: A Parameter-free Deep Learning Framework for Nucleus Segmentation Using Image Style Transfer. Cell Syst. (2020), doi:10.1016/j.cels.2020.04.003.

40. Y. Xue, S. Cheng, Y. Li, L. Tian, Reliable deep-learning-based phase imaging with uncertainty quantification. Optica. 6, 618–629 (2019).

41. R. Liu, S. Cheng, L. Tian, J. Yi, Deep spectral learning for label-free optical imaging oximetry with uncertainty quantification. Light Sci. Appl. 8, 102 (2019).

42. A. Krull, T.-O. Buchholz, F. Jug, in Proceedings of the IEEE Conference on Computer Vision and Pattern Recognition (2019), pp. 2129–2137.

43. S. Li, M. Deng, J. Lee, A. Sinha, G. Barbastathis, Imaging through glass diffusers using densely connected convolutional networks. Optica. 5, 803–813 (2018).

44. N. Otsu, A Threshold Selection Method from Gray-Level Histograms, 5.

45. A. E. Carpenter, T. R. Jones, M. R. Lamprecht, C. Clarke, I. H. Kang, O. Friman, D. A. Guertin, J. H. Chang, R. A. Lindquist, J. Moffat, P. Golland, D. M. Sabatini, CellProfiler: image analysis software for identifying and quantifying cell phenotypes. Genome Biol. 7, R100 (2006).

