## Supplementary Materials for "Single-cell cytometry via multiplexed fluorescence prediction by label-free reflectance microscopy"

Section S1. Outlier detection and removal
Section S2. Procedure of quantifying cell-level detection performance
Section S3. Procedure of digital cytometry analysis
Section S4. Cell profile analysis
Section S5. Computation of the saliency map

Fig. S1. Neural network structure
Fig. S2. Data preprocessing pipeline
Fig. S3. Network prediction with and without drDPC input channels
Fig. S4. Additional individual prediction results
Fig. S5. Spatial variation quantification of prediction accuracy
Fig. S6. Additional multiplexed prediction results
Fig. S7. Additional single-cell profile metrics
Fig. S8. Procedure of quantifying cell-level detection performance
Fig. S9. Procedure of digital cytometry analysis

Table S1. Quantification of network prediction with and without drDPC input channels

### Supplementary Materials

#### Section S1. Outlier detection and removal

When computing the patch-wise PCC, we noticed that the lower accuracy regions generally align with the background region, where the local signal density and SNR is relatively low. This means that our patch-wise PCC computation is sensitive to low-SNR regions and PCC outliers in these regions should be eliminated. We added a median-based outlier detection and removal algorithm to remove the outliers and all the sub-FOV PCC shown on our violin plots in Fig. 3. The outlier detection and removal algorithm are implemented by a Matlab built-in function '`filloutliers`' with `findmethod` of '`median`' and `fillmethod` of '`pchip`'. The '`median`' method finds the outliers as values that are three scaled median absolute deviations (MAD) away from the median. The '`pchip`' method fills the outliers using shape-preserving piecewise cubic spline interpolation.

**Section S2. Procedure of quantifying cell-level detection performance**

To investigate the effect of false predictions on the generated digital cytometry scatter plot, we develop an image processing pipeline using CellProfiler to quantify the cell-level performance. For both proliferation and apoptosis, the TP, TN, FP, and FN cells are individually identified by generating the corresponding nuclei masks using the same pipeline. Those image masks are stored in separate files and counted separately. The procedure of segmenting TP/TN/FP/FN nuclei is described below, and shown in Fig. S8.

**1)** To generate TP nuclei masks, the ground truth labeled nuclei are first segmented, which are denoted by GTP, using '`IdentifyPrimaryObjects`' module.

**2)** The predicted positively labeled nuclei are segmented using the same module, denoted as POS.

**3)** We then utilized '`RelateObjects`' module to label POS with GTP as reference and '`FilterObjects`' module to filter out POS that *does not contain* GTP labels. This operation takes the intersection of POS and GTP, which are TP nuclei denoted by TP here.

**4)** We then label POS with TP as the reference and filter out POS that *contains* TP labels using the same modules. This operation takes the complement of TP from POS to get the FP nuclei, denoted as FP.

**5)** The co-stained DNA channel ground truth are then used to segment all nuclei, denoted as ALL.

**6)** We used the same complement operation on ALL with reference POS to get NEG and on ALL with reference GTP to get GTN.

**7)** Then we intersect NEG with GN using the same operation as described in (3) to pick out true negatives TN.

**8)** False negatives FN are then obtained by taking the complement NEG with TN reference using the same operation as described in (4).

For proliferation fluorescence prediction, the intensity values are also aggregated within each type of cell mask regions to mark the scatter plot with individual detection performance labels. The segmentation and measurement procedure is the same as that described in Section S3.

### Section S3. Procedure of digital cytometry analysis

To generate the cytometry scatter plots, we feed the co-registered Hoechst and EdU images to CellProfiler and then perform the following steps and shown in Fig. S9.

We segment all the nuclei in the Hoechst image by the '`IdentifyPrimaryObjects`' module and extract all the corresponding nuclei masks in the data. We then apply the masks to both Hoechst and EdU images and calculate the integrated fluorescence intensity inside the mask by the '`MeasureObjectIntensity`' module.

One confounding factor of this pipeline we find is that using the pure image data can mis-detect certain M-phase cells during the segmentation. Specifically, in anaphase and telophase (sub-phases in M phase) when a cell is about to split into two, the typical morphology is that a cell is attaching or intimately close to a nearby one. While common flow cytometry readout treats the two cells as one (since they are likely to be flown through the fluidic channel simultaneously) and gives the correct readings of DNA and proliferating DNA expression, our digital cytometry does not with a plain segmentation algorithm in CellProfiler. To overcome this issue, we refine the segmentation procedure by the following steps.

**1)** We set the segmenting size to 24-35 pixels to find normal-sized nuclei that are prevalent in S and G1/G2 phases.

**2)** We set the segmenting size to 16-23 pixels to find small-sized nuclei that exist in the M phase.

**3)** We use the '`MergeObjects`' module to combine the small-sized nuclei masks that are within a distance of 2 pixels into a single mask.

**4)** We combine the merged masks and the normal-sized nuclei masks for computing the cell-level fluorescence intensities.

We find this procedure successfully reduces the misdetection of M-phase nuclei and provides a more accurate scatter plot.

**5)** The paired cell-level fluorescence intensity data are then plotted in Matlab. A small value ($10^{-10}$) is added to the fluorescence intensity of the proliferating DNA before taking the log operation to avoid singularity at 0.

For the multiplexed predictions, we apply the same pipeline on the predictions from the DNA and proliferation networks. Three distinct clusters representing the S, G1, and G2/M phase are clearly shown in the digital cytometry scatter plots.

### Section S4. Cell profile analysis

We use CellProfiler to generate the single-cell profiles across each entire fluorescence image. We feed the ground truth and the predicted IF images of DNA, actin, endosome and Golgi apparatus to CellProfiler. After initial cell segmentation, single-cell level parameters of morphology and intensity distribution are computed automatically by different measurement modules in CellProfiler. Specifically, we use the '`MeasureObjectSizeShape`' module to measure the fluorescence marker size (area), compactness, and eccentricity, the '`MeasureObjectIntensity`' module to measure the fluorescence concentration, the '`MeasureTexture`' module to measure the single-cell-level fluorescence variance and contrast. The fluorescence marker size is measured by the number of pixels in the segmented cell region. The compactness is computed by the mean squared distance of the cell mask pixels from the centroid divided by the area. The eccentricity is defined by the ratio of the distance between the foci of the effective ellipse that has the same second-moments as the segmented region and its major axis length. The concentration is computed as the sum of the intensities within the segmented masks. The contrast is measured by the local variation in a cell region. The variance is measured by the variation of the intensity values. The cell-level parameters are extracted from CellProfiler and then imported to Matlab to generate the violin plots. NCR is a compound metric that involves multiplexed DNA and actin fluorescence labels. To compute this metric, co-registered ground-truth / prediction images containing the DNA and actin labels are processed individually in CellProfiler. NCR is computed as the ratio between the area of segmented nuclei and actin masks.

### Section S5. Computation of the saliency map

We utilized a built-in function ‘`visualize_saliency`’ in ‘`vis.visualization`’ module from an open-source packages ‘`Keras-vis`’ to enable the gradient computation (*41*). This is conveniently achieved by the following single-line script with several arguments to be specified.

```
saliency_map= visualize_saliency(model, layer_idx=137,
seed_input=seed, grad_modifier='absolute',
backprop_modifier='guided')
```

Our output layer is specified as the last layer - layer 137 with the seed or input being the first layer of the network. We specified the gradient modifier as absolute values of the gradient, which shows the regions in the input that contribute most to the change in the output regardless of the sign of the change (i.e. negative or positive). We also used the guided backpropagation to propagate only the positive gradients for positive activations to achieve a smoother gradient visualization.

**Figures and Tables**

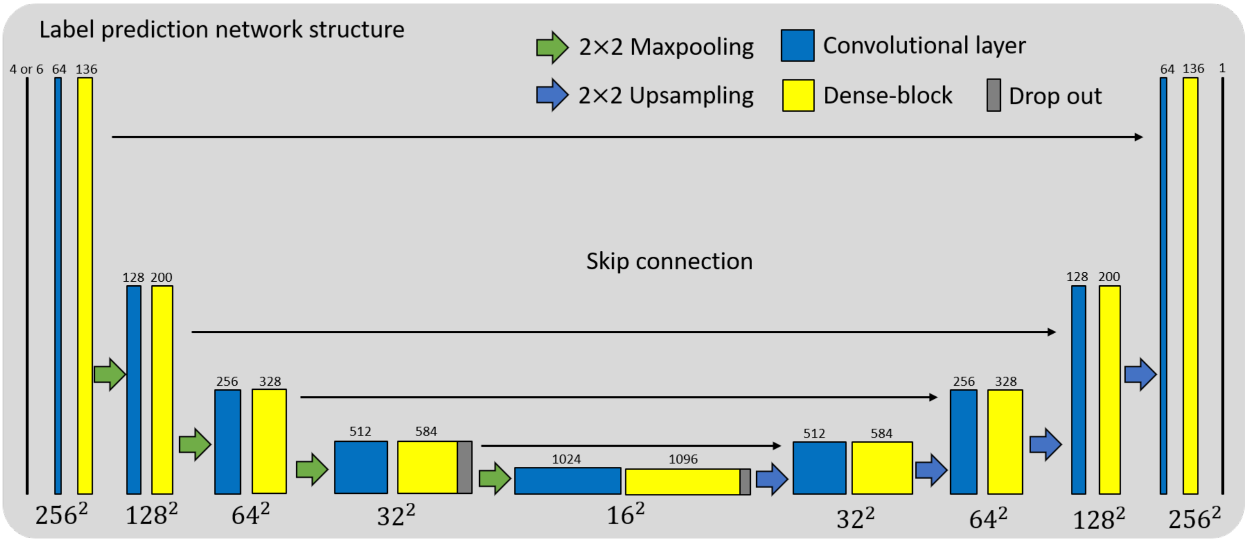


**Fig. S1. Neural network structure.** The overall structure of the neural network follows the U-Net architecture modified with the dense-block modules. The input takes 256×256-pixel label-free images consisting of four darkfield reflectance images. For predicting the actin fluorescence labels, the input is expanded to six channels that include additionally two-directional drDPC images. For all the conditions, the output predicts a single-channel fluorescence image with 256 × 256 pixels. Starting with the multi-channel high-resolution label-free reflectance images, the encoder path gradually condenses the lateral spatial information (size marked by the number in the bottom of each block) into high-level feature maps with growing depths (size marked by the number in the top of each block); the decoder path reverses the process by recombining the information into feature maps with gradually increased lateral details. The information in adjacent feature maps (painted in blue and yellow) transfers by convolving with 3 × 3 convolutional filters. The downsampling is done by 2 × 2 maxpooling operation and the upsampling is done by 2 × 2 upsampling operation. Dense-blocks are added to facilitate efficient training. Each dense-block contains multiple layers, in which each layer consists of batch normalization (BN), the rectified linear unit (ReLU) nonlinear activation, and convolution (conv) with 24 filters. We also added drop out layers with 0.5 dropout rate after the central two dense-blocks to mitigate overfitting. Skip connections are added to tunnel the high-frequency information from shallower layers to deeper layers with the same spatial scales. We use ReLU for all activation functions of the hidden layers and the sigmoid activation function in the final layer to form the final prediction ranging between 0 and 1.

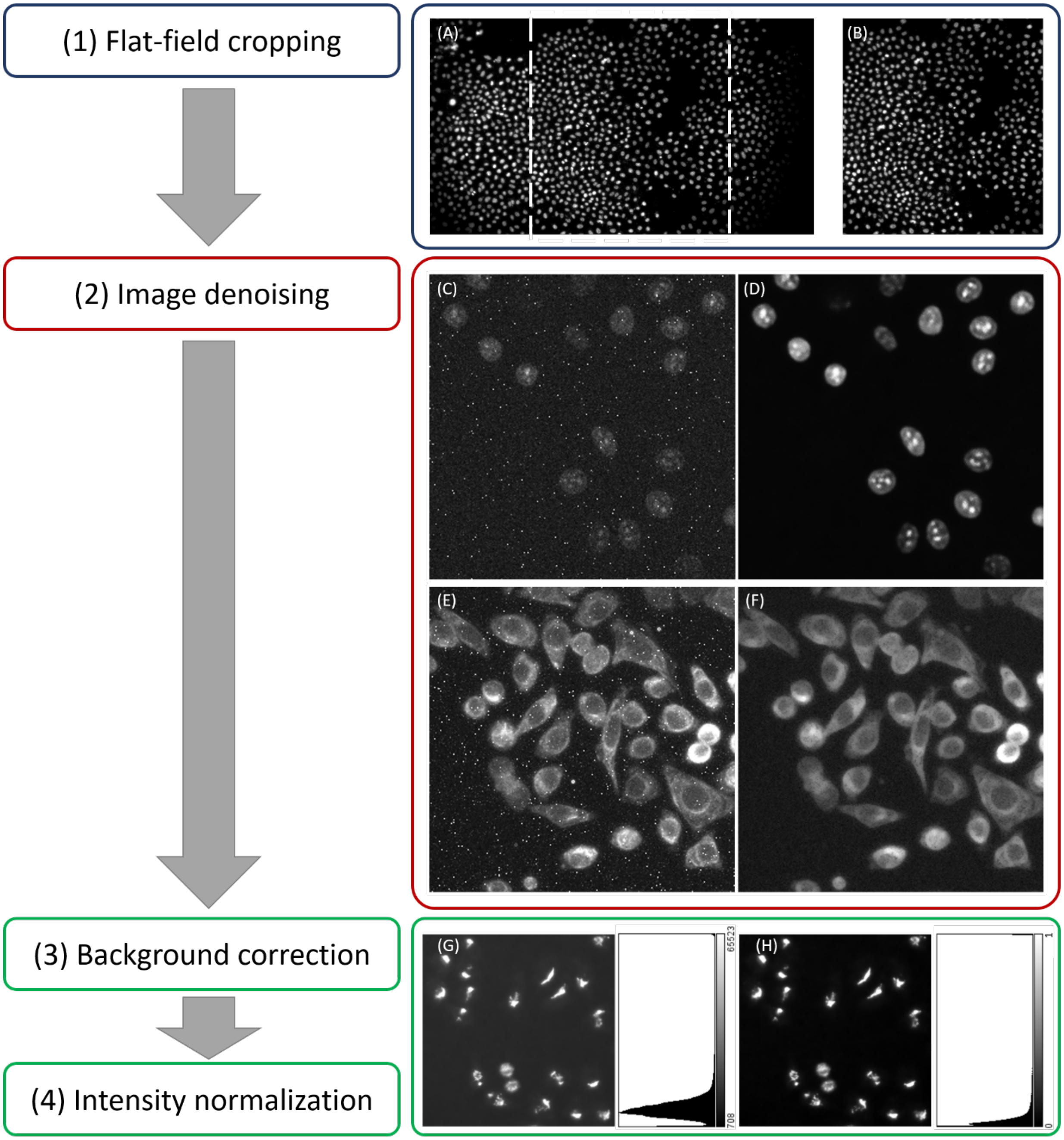


**Fig. S2. Data preprocessing pipeline.** (**A**) An example of Hoechst stained DNA image with the whole 1920×1080-pixel FOV. The image suffers from non-uniform fluorescence expression due to non-uniform excitation illumination. (**B**) The image is processed by the flat-field cropping step that maintains only the central 1000×1080-pixel region that retains approximately a uniform illumination. (**C**) An example of low-SNR raw measurement of the proliferation label that is contaminated by hot pixels and sensor noise. (**D**) The denoised proliferation-label image by the noise2void unsupervised deep learning algorithm. Both the sensor noise and the hot pixels are removed while maintaining the fluorescence signals with high fidelity. (**E**) An example of raw measurement of the endosome label that is primarily contaminated by hot pixels and with relatively low sensor noise. (**F**) The hot pixels are removed by morphological opening operation. (**G**) An example image and the histogram of measurement for the Golgi apparatus label that contains a constant background offset. (**H**) The processed Golgi label image and its histogram with the corrected background by subtracting the offset and normalized intensity range.

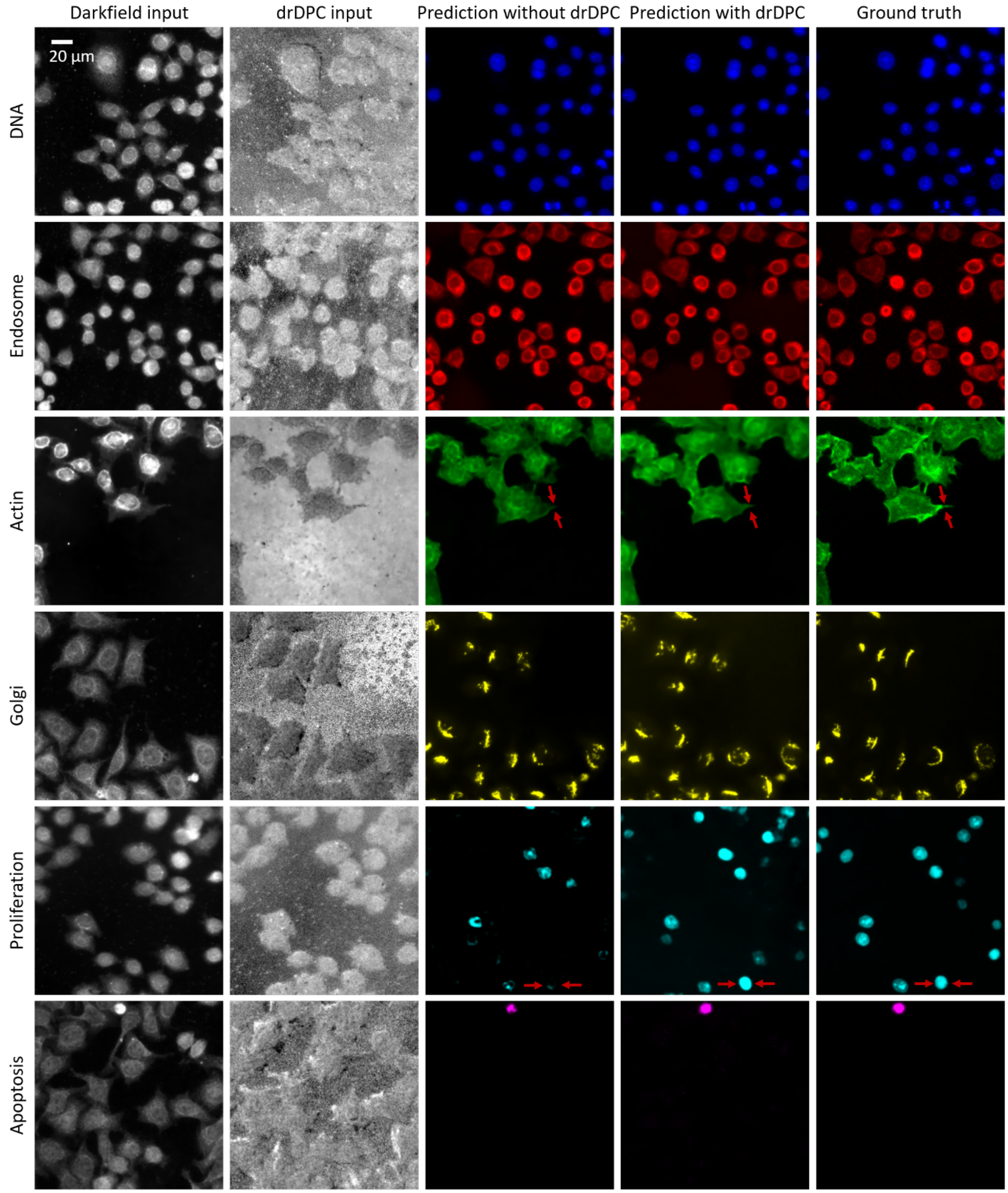


**Fig. S3. Network prediction with and without drDPC input channels.** Example prediction results of six labelling networks trained with two different input settings: one with only four-channel darkfield reflectance input, the other with four-channel darkfield reflectance and two additional drDPC images as the input. The first two columns show examples of the darkfield reflectance and drDPC images used for DNA, endosome, actin, Golgi, proliferation and apoptosis prediction, respectively. The last three columns show the predicted labels without the drDPC input, with the drDPC input and the ground truth labels, respectively. The drDPC input helps preserving the fine details in the actin boundaries and improving true positive predictions in proliferation label, as shown in the examples marked by the red arrows.

| Fluorescence labels | DNA | Endosome | Actin | Golgi | Proliferation | Apoptosis |
|---|---|---|---|---|---|---|
| Accuracy without drDPC | **87.25%** | **91.85%** | 83.25% | **59.82%** | 83.90% | 60.20% |
| Accuracy with drDPC | 85.20% | 90.07% | **92.01%** | 57.20% | **91.65%** | **77.24%** |

**Table S1. Quantification of network prediction with and without drDPC input channels.** Quantitative metrics including PCC and AUC on 128 testing set images for six fluorescence predictions from models trained with two different input settings, including four-channel plain darkfield images of different illumination patterns or six-channel images consisting of the four-channel darkfield images and additional two-channel drDPC images. The prediction accuracy of DNA, endosome, Golgi degraded slightly with the additional drDPC input channels possibly because the drDPC morphological features do not add useful information their predictions. The performance substantially improved for actin, proliferation and apoptosis with the additional drDPC input channels, indicating that the DPC morphological features contain rich information for their predictions.

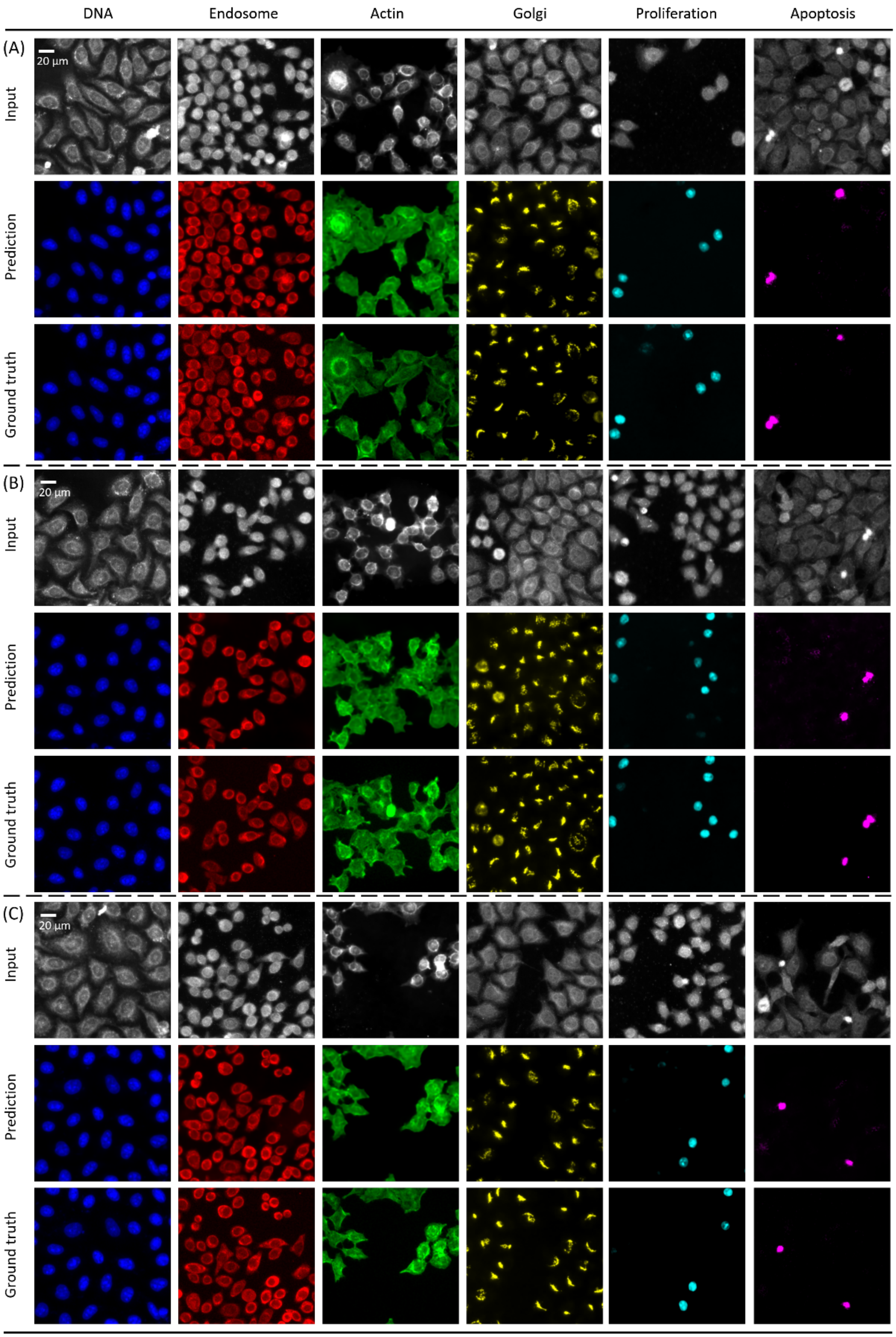


**Fig. S4. Additional individual prediction results. (A-C)** The rows show a sample reflectance image from each input stack, the network's prediction, and the ground-truth IF image, respectively. The columns represent six IF labels covering four different subcellular features, including nuclei (DNA), endosome, actin, and Golgi apparatus, as well as two different cell states, including proliferation and apoptosis.

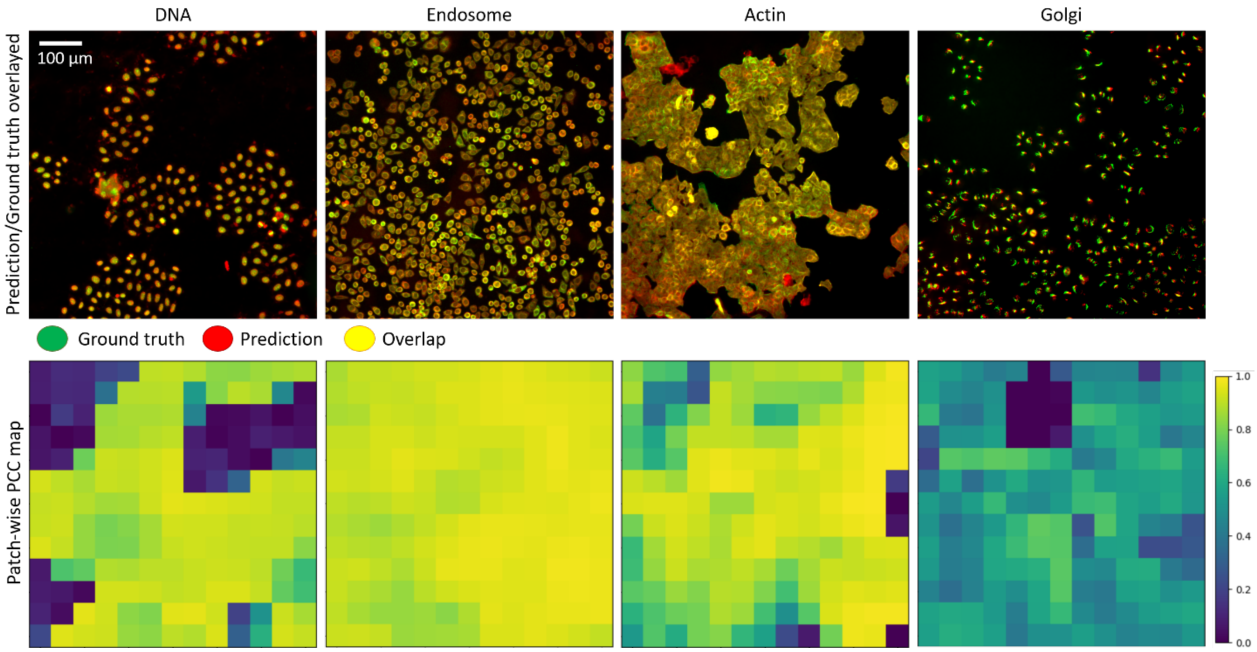


**Fig. S5. Spatial variation quantification of prediction accuracy.** The prediction and ground truth images across 908×908-pixel FOVs along with their patch-wise PCC map displayed on a 13×13 grid. Each PCC value is computed from a 128×128-pixel image patch; the map is computed with a 64×64-pixel overlap between the neighboring image patches. (Top row) Red: the prediction of four 'regression' labels, including the DNA, endosome, actin, Golgi apparatus; Green: the ground truth overlay. (Bottom row) the corresponding PCC maps. The low PCC regions show spatial correspondence with the background region in the sample. Other than the background outliers, the PCCs are consistent across all the cell regions for all the label predictions.

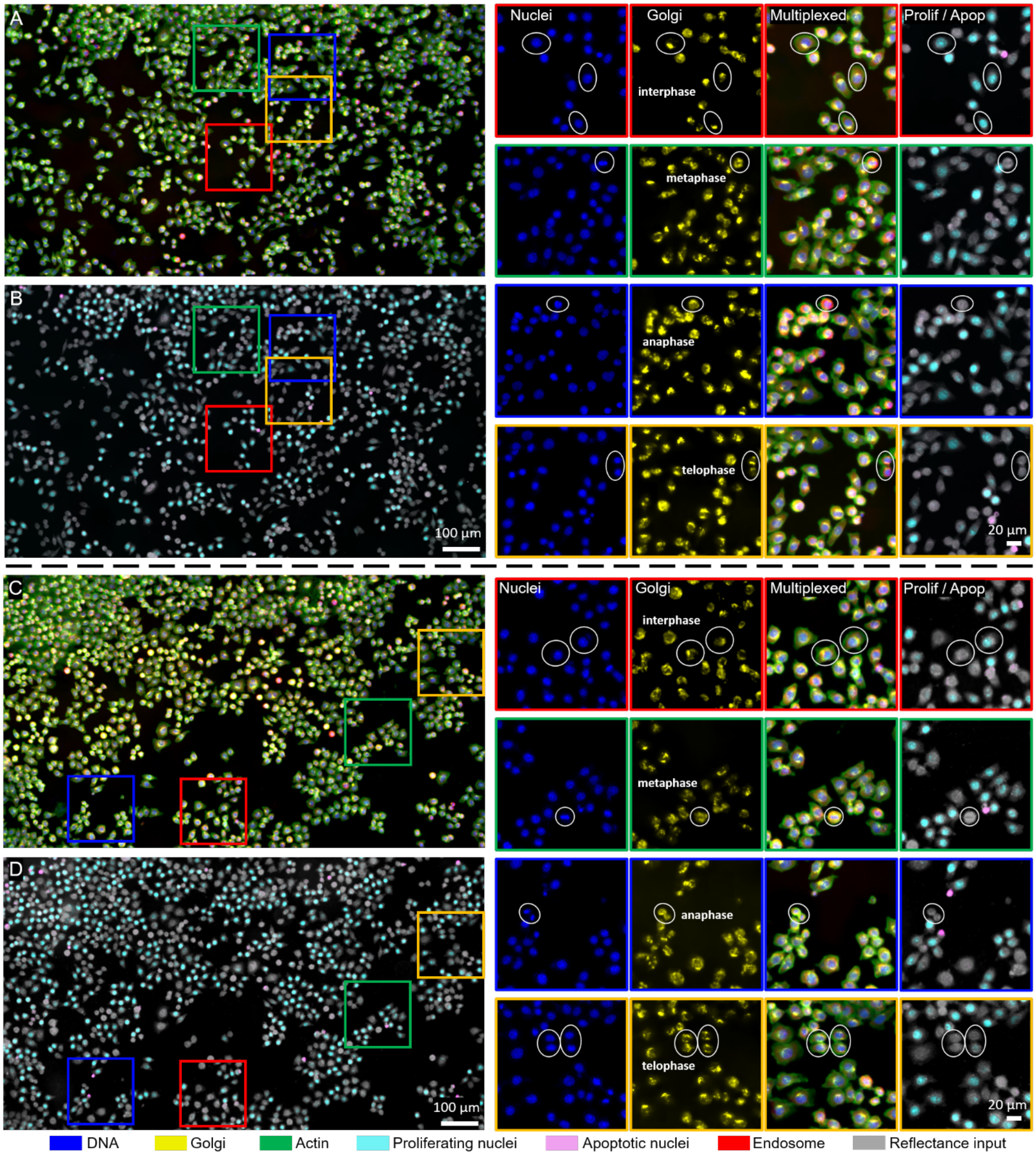


**Fig. S6. Additional multiplexed prediction results. (A, C)** Visualization of the Full-FOV multiplexed prediction including DNA (blue), endosome (red), actin (green), and Golgi apparatus (yellow), and **(B, D)** proliferation (cyan) and apoptosis (magenta) from the same reflectance input (grayscale). The right-hand-side are zoomed-in of DNA, Golgi apparatus, multiplexed, and proliferation predictions. White circles indicate representative cell morphology during different phases of cell cycle, including interphase, metaphase, anaphase and telophase. **(A, B)** Multiplexed predictions from the endosome staining cell batch. **(C, D)** Multiplexed predictions from the proliferation staining cell batch.

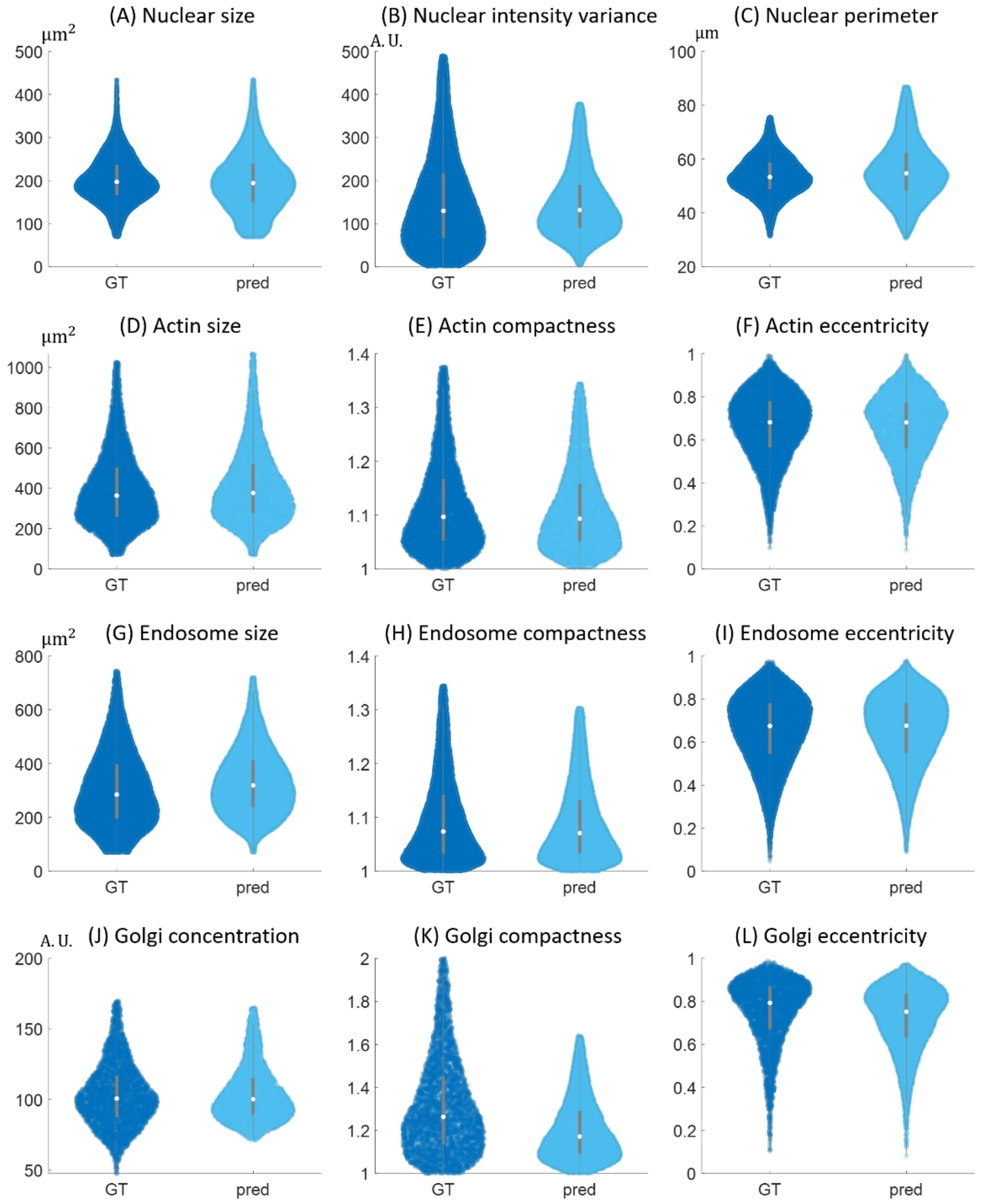


**Fig. S7. Additional single-cell profile metrics.** The comparisons of the statistics of twelve different single-cell profile metrics extracted from the entire cell population in the ground truth (GT) and DL-predictions (pred), including **(A)** the nuclear size, **(B)** the DNA (nuclear) fluorescence intensity variance, **(C)** the nuclear perimeter, **(D)** the cell (actin) size, **(E)** the compactness of actin, **(F)** the eccentricity of actin, **(G)** the endosome size, **(H)** the compactness of endosome, **(I)** the eccentricity of endosome, **(J)** the concentration of Golgi apparatus, **(K)** the compactness of Golgi apparatus distribution, and **(L)** the eccentricity of Golgi apparatus distribution. The total numbers of sample cells collected from ground truth and DL-predictions are respectively: 20021 and 25257 in plots **(A) (B) (C)**, 6183 and 5151 in plots **(D) (E) (F)**, 15944 and 22408 in plot **(G) (H) (I)**, 3380 and 9392 in plot **(J) (K) (L)**. All the single-cell profile metrics show good agreements between the prediction and the ground truth.

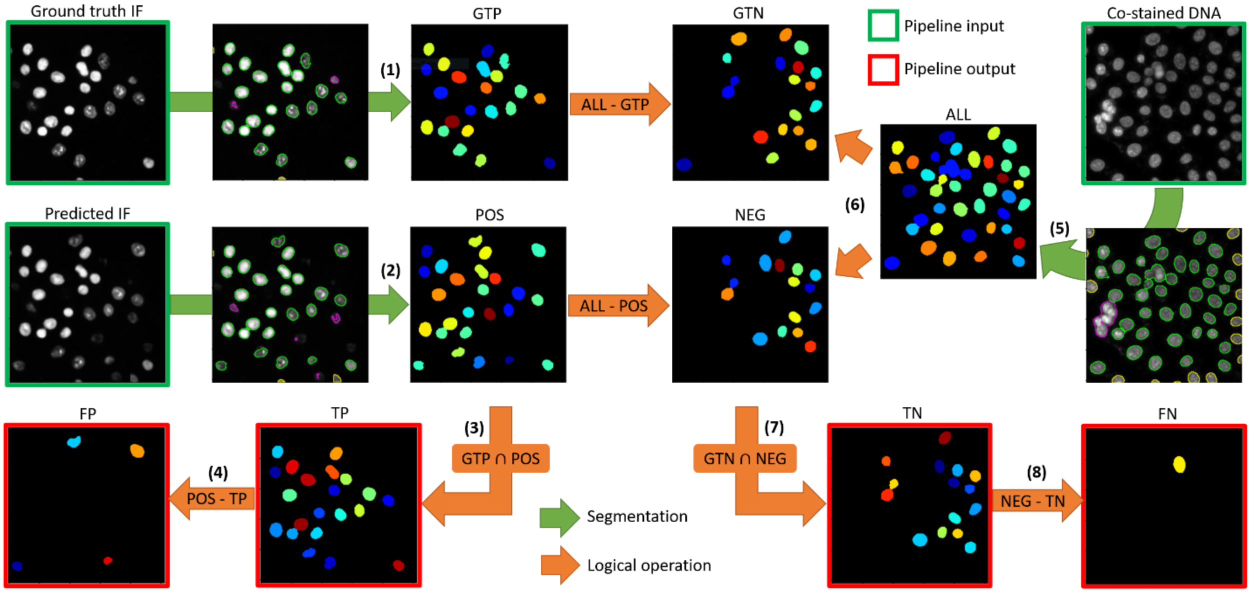


**Fig. S8. Procedure of quantifying cell-level detection performance.** Image processing pipeline for the quantification of four cell-level detection outcomes, including true positives (TP), false positives (FP), true negative (TN) and false negatives (FN). All the detection performances are in terms of the concentration of target IF. The segmentation and logical operation are implemented by the modules in CellProfiler as described in Section S2.

**(1)** Ground truth target IF images are fed into the segmentation module to locate the ground truth positive (GTP) cells.

**(2)** Predicted target IF images are segmented to find the predicted-positive (POS) cells.

**(3)** The intersecting operation on GTP and POS provides the true positives (TP).

**(4)** Taking the complement of TP from POS yields the false positives (FP).

**(5)** The co-stained DNA channel is segmented to locate all (ALL) the cells.

**(6)** The complement of GTP from ALL is the ground truth negatives (GTN); the complement of POS from ALL is the predicted-negatives (NEG).

**(7)** The intersection between GTN and NEG is TN.

**(8)** The complement of TN from NEG is FN.

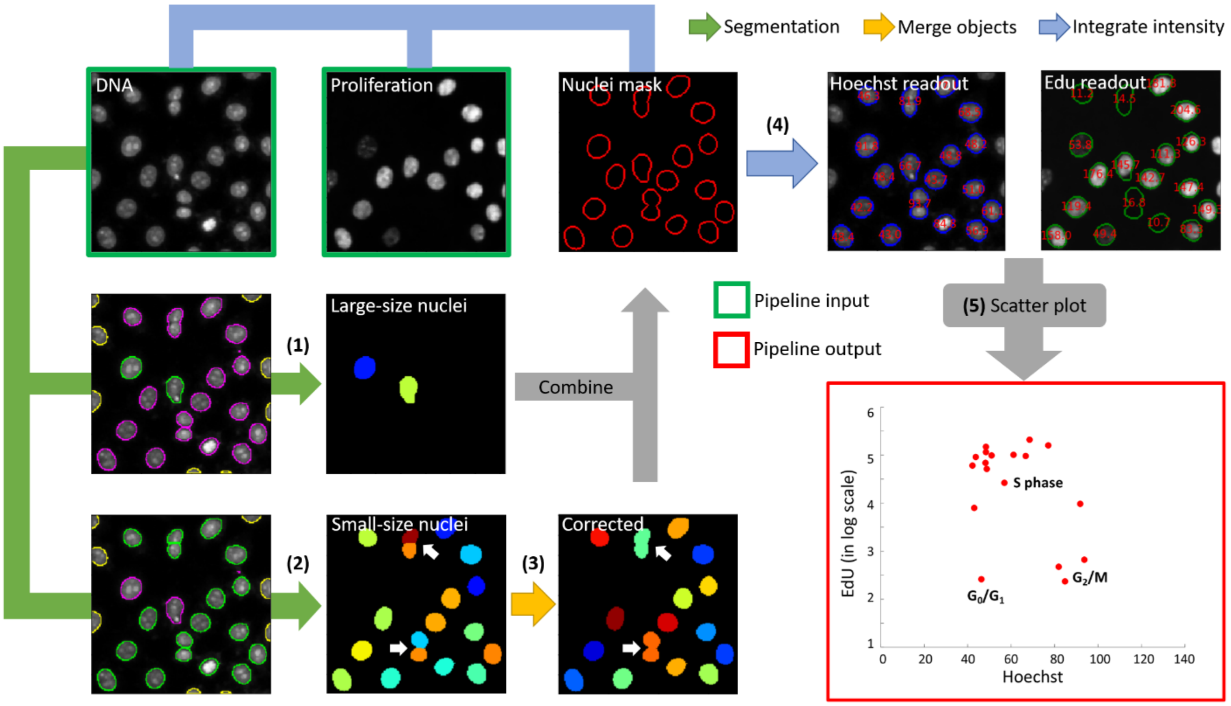


**Fig. S9. Procedure of digital cytometry analysis.** Image processing pipeline for digital cytometry scatter plot. The pipeline applies to the DNA/proliferation image pairs from both the ground truth and the multiplexed predictions. The segmentation, 'merge objects' and 'integrate intensity' operations are implemented by the modules in CellProfiler as described in Section S3.

**(1)** The DNA images are segmented to locate all the large-size nuclei using a large size constraint.

**(2)** The same DNA images are then segmented to find all the small-size nuclei with a smaller size constraint.

**(3)** An additional correction operation is performed on small-size nuclei to merge the incorrectly separated cell segments (those during mitosis) into a single segment. The corrected small-size nuclei masks are then combined with the large-size masks to provide a union of all the nuclei masks.

**(4)** The DNA and proliferation image pairs are then masked. The intensity values within each mask are aggregated to provide the single cell-level concentration readout. The DNA channel corresponds to the Hoechst-stain concentration and the proliferation channel correspond to the Edu-stain concentration.

**(5)** All the single-cell level two-channel readouts are plotted cell by cell.
